# Mitotic chromosome condensation requires phosphorylation of the centromeric protein KNL-2 in *C. elegans*

**DOI:** 10.1101/2021.07.01.450752

**Authors:** Joanna M. Wenda, Reinier F. Prosée, Caroline Gabus, Florian A. Steiner

**Affiliations:** Department of Molecular Biology and Institute for Genetics and Genomics in Geneva, Section of Biology, Faculty of Sciences, University of Geneva, 1211 Geneva, Switzerland

**Keywords:** KNL-2, chromosome condensation, *C. elegans*, condensin II, centromere

## Abstract

Centromeres are chromosomal regions that serve as sites for kinetochore formation and microtubule attachment, processes that are essential for chromosome segregation during mitosis. Centromeres are almost universally defined by the histone variant CENP-A. In the holocentric nematode *C. elegans*, CENP-A deposition depends on the loading factor KNL-2. Depletion of either CENP-A or KNL-2 results in defects in centromere maintenance, chromosome condensation and kinetochore formation, leading to chromosome segregation failure. Here, we show that KNL-2 is phosphorylated by CDK-1, and that mutation of three C-terminal phosphorylation sites causes chromosome segregation defects and an increase in embryonic lethality. In strains expressing phosphodeficient KNL-2, CENP-A and kinetochore proteins are properly localised, indicating that the role of KNL-2 in centromere maintenance is not affected. Instead, the mutant embryos exhibit reduced mitotic levels of condensin II on chromosomes and significant chromosome condensation impairment. Our findings separate the functions of KNL-2 in CENP-A loading and chromosome condensation and demonstrate that KNL-2 phosphorylation regulates the cooperation between centromeric regions and the condensation machinery in *C. elegans*.

**SUMMARY STATEMENT:** Phosphorylation of the essential centromere protein KNL-2 is required for mitotic chromosome condensation, but not for the role of KNL-2 in centromere maintenance and kinetochore formation.

## INTRODUCTION

At the onset of mitosis, the loose interphase chromatin condenses into compact, rod-shaped chromosomes that are subsequently pulled apart by the mitotic spindle to the forming daughter cells. The points of contact between the spindle and chromosomes are special regions of chromatin, called centromeres, that are exposed on the surface of mitotic chromosomes and positioned to face opposite sides. Centromeres have a specific structural organisation enabling them to withstand forces exerted by the spindle. They recruit kinetochore complexes, multiprotein structures that mechanically anchor the spindle microtubules onto the chromatin (McKinley and Cheeseman, 2016). The specific functions of centromeres require properties and adaptations that are different from non-centromeric chromatin (Bloom and Joglekar, 2010).

Despite sharing a common principle of action, centromeres vary in organisation between different species. In monocentric species, centromeres occupy a restricted region of the chromosome with sizes ranging from 125 base pairs in budding yeast to millions of base pairs in vertebrates (McKinley and Cheeseman, 2016). In holocentric species, centromeres cover the whole axis of the chromosome (Steiner and Henikoff, 2015). The defining feature for centromeres in most eukaryotes is the presence of CENP-A, a centromeric variant of histone H3 (McKinley and Cheeseman, 2016).

Although the maintenance of centromeric chromatin mechanistically differs between species, it generally revolves around the timely deposition of CENP-A. Licensing factors, such as the MIS18 complex (Fujita et al., 2007; Hayashi et al., 2004) identify the sites for CENP-A deposition and recruit CENP-A specific chaperones, HJURP or Scm3 (Bernad et al., 2011; Dunleavy et al., 2009; Foltz et al., 2009), which then complete the process of loading of new CENP-A nucleosomes. CENP-A loading is often restricted to a particular phase of the cell cycle, e.g. G1 in human cells (Jansen et al., 2007) or G2 in *Schizosaccharomyces pombe* (Dunleavy et al., 2007). The spatiotemporal regulation of CENP-A deposition is heavily dependent on phosphorylation events carried out by mitotic kinases. In human cells, the CENP-A licensing machinery is inhibited by CDK1-mediated phosphorylation of the MIS18 licensing complex and HJURP (McKinley and Cheeseman, 2014; Pan et al., 2017; Silva et al., 2012; Spiller et al., 2017; Stankovic et al., 2017). PLK1 phosphorylates the MIS18 complex to promote CENP-A deposition (McKinley and Cheeseman, 2014). The recruitment of inner and outer kinetochore proteins is also regulated by phosphorylation events (Navarro and Cheeseman, 2021).

The spatial organisation of centromeres within mitotic chromosomes is important for their specific mechanistic properties. Centromeres are typically denser than the rest of the chromosome and more resistant to the tension created by the mitotic spindle pulling forces (Bloom and Joglekar, 2010; Harasymiw et al., 2019). Centromere elasticity and tension-sensing mechanisms are thought to play an important role in achieving bi-orientation and faithful chromosome segregation (Foley and Kapoor, 2013). Furthermore, condensin complexes involved in the formation of the mitotic chromosomes (Gibcus et al., 2018; Hirano and Mitchison, 1994; Ono et al., 2003) are enriched at centromeric regions (Csankovszki et al., 2009; Hagstrom et al., 2002; Ono et al., 2004; Savvidou et al., 2005; Shintomi and Hirano, 2011) and influence centromeric chromatin organisation (Bernad et al., 2011; Oliveira et al., 2005; Samoshkin et al., 2009; Yong-Gonzalez et al., 2007).

Holocentric chromosomes provide an especially interesting model for studying centromere establishment and the properties of centromeric regions. Centromeric chromatin is not restricted to a specific region, but scattered discontinuously across the genome, which may require special adaptations for its maintenance machinery. Furthermore, on the condensed chromosomes, these scattered regions are all placed on the surface and collectively span the whole chromosome axis (Steiner and Henikoff, 2015). This centromere organisation might have unique consequences for chromosome condensation and their physical properties.

In the holocentric nematode *Caenorhabditis elegans* the functional CENP-A homologue is called HCP-3 (thereafter referred to as CENP-A for clarity) (Buchwitz et al., 1999; Monen et al., 2005). CENP-A deposition is regulated by KNL-2, a M18BP1 homologue, which acts as a loading factor in *C. elegans* (Maddox et al., 2007), and LIN-53, a RbAp46/48 homologue (Lee et al., 2016). KNL-2 and CENP-A interact directly through CENP-A N-terminal tail and are dependent on one another for chromatin binding (de Groot et al., 2021; Maddox et al., 2007; Prosée et al., 2020). They exhibit a similar localisation pattern throughout the cell cycle and have overlapping genomic distributions (Gassmann et al., 2012; Maddox et al., 2007). On prometaphase chromosomes they localise to form a characteristic pattern of two parallel lines (‘railroad track’) spanning the entire chromosome length. CENP-A and KNL-2 are required for kinetochore recruitment (Maddox et al., 2007; Oegema et al., 2001). Depletion of either KNL-2 or CENP-A is detrimental for cell viability and results in severe cell division impairment: chromosomes do not condense properly and the kinetochores fail to assemble leading to lack of microtubule attachment and a failure in chromosome segregation (Hagstrom et al., 2002; Maddox et al., 2007, 2006; Oegema et al., 2001).

However, it remains unclear if all the mitotic defects observed after KNL-2 depletion are a consequence of the failure to load CENP-A and form centromeres. Chromosome condensation and segregation are dynamic processes that are mechanistically linked, and secondary defects in depletion experiments could obscure important roles of KNL-2 in either of these processes. To investigate the roles of KNL-2 in more detail, we therefore examined KNL-2 post-translational modifications that could be involved in its spatiotemporal regulation.

Here we show that the function of KNL-2 in mitosis is regulated by phosphorylation in *C. elegans* embryos. Mutation of three CDK-1 phosphorylation sites results in cell division defects and embryonic lethality. While CENP-A loading and kinetochore recruitment are not affected in the phosphodeficient strain, chromosome condensation is significantly impaired. These observations show that the KNL-2 functions in chromosome condensation and centromere maintenance are independent and separately regulated. We propose that KNL-2 is a main player in orchestrating the cooperation between centromeric chromatin and the condensation machinery in *C. elegans* embryos.

## RESULTS

### KNL-2 is regulated by phosphorylation

Depletion of KNL-2 from *C. elegans* embryos results in severe mitotic defects, including loss of CENP-A on chromatin, inability to form the kinetochore, defects in chromosome condensation, and failure to segregate chromosomes (Maddox et al., 2007). In order to investigate how KNL-2 function in these processes is regulated, we set out to identify potential regulatory post-translational modifications. We created a strain expressing HA-tagged KNL-2 from the endogenous locus and performed immunoprecipitation from worm embryonic lysates followed by mass spectrometry. We focused our analysis on phosphorylated residues fitting the motif S/TP, the minimal consensus for cyclin-dependent kinases (CDKs) (Errico et al., 2010), since centromere licensing factors in other species are regulated by CDK-1 phosphorylation (French and Straight, 2019; McKinley and Cheeseman, 2014; Pan et al., 2017; Silva et al., 2012; Spiller et al., 2017; Stankovic et al., 2017). We identified two such sites near the C-terminus of KNL-2 (Fig. 1A, Fig. S1A), S772 and S784. Another candidate CDK target TP site (T750) is present in close proximity. We did not find evidence for its phosphorylation in our mass spectrometry data, possibly because the peptide generated after trypsin digestion would be very short (Table S1). All three sites are conserved within the *Caenorhabditis* genus (Fig. S1B), and we hypothesised that all three could serve as targets for phosphorylation for the cyclin-dependent kinase CDK-1.

**Figure 1.**
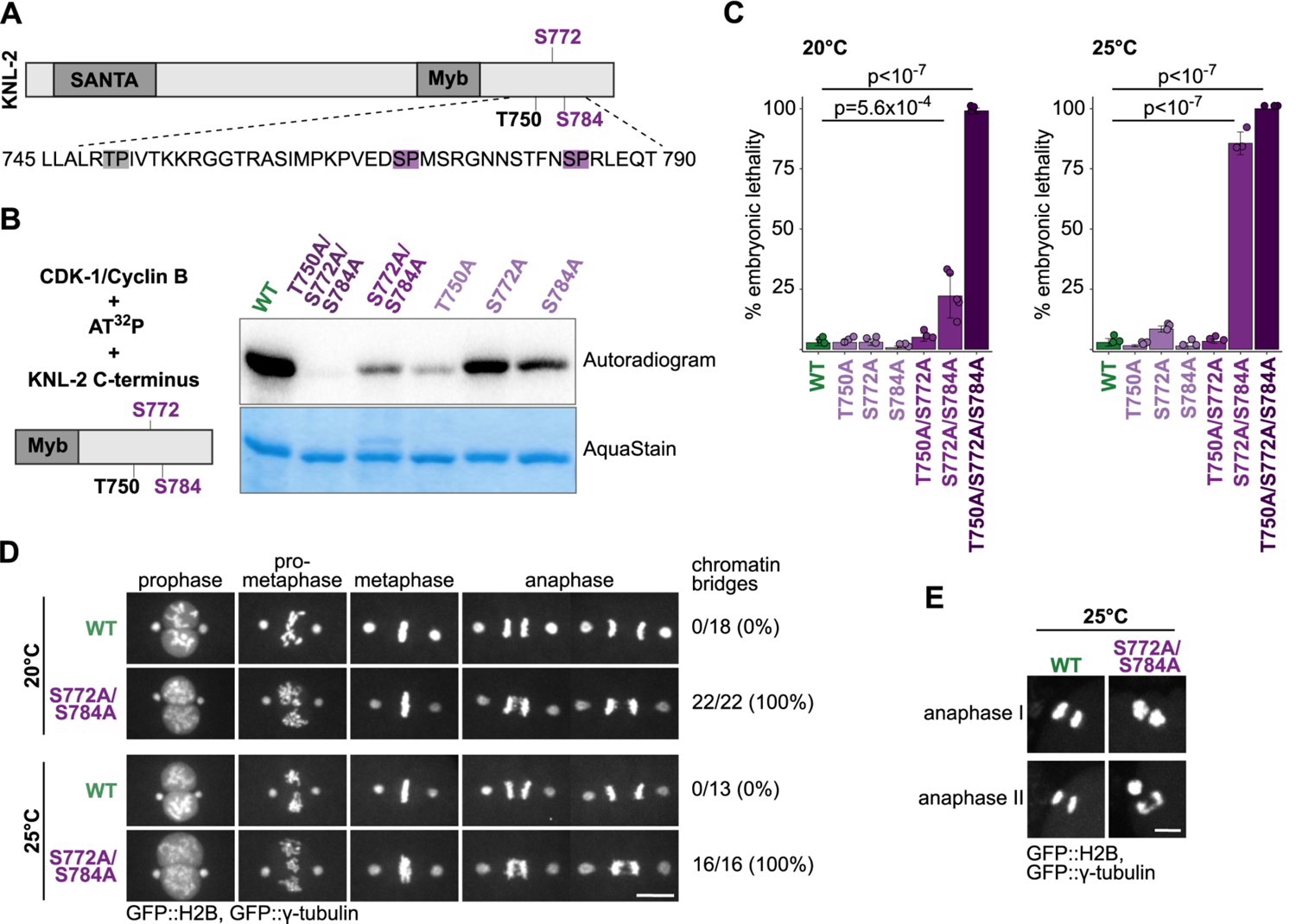
The C-terminal part of KNL-2 is phosphorylated by CDK-1. (A) Top, scheme of KNL-2 with annotated domains and phosphosites analysed in this study. Bottom, amino acid sequence containing the three phosphosites with CDK-1 consensus sequence. Two phosphosites identified by the IP-MS analysis are marked in violet, a third site with a CDK-1 consensus sequence is marked in grey. (B) *In vitro* CDK-1 kinase assay for recombinant C-terminal fragments of KNL-2, with the phosphosites mutated to alanines individually or in combination with each other. Left, scheme of the reagents used. Right, autoradiogram and AquaStain staining of the purified KNL-2 fragments. (C) Quantification of embryonic lethality for wild type (WT) and different KNL-2 phosphosite mutants at 20°C or 25°C. Data points indicate the average value for each biological replicate of the experiment (8-10 hermaphrodites used and a total of over 100 embryos were scored per replicate). For statistical analysis ANOVA followed by Tukey-Kramer post hoc was used. Relevant p-values are indicated, all other comparisons to WT are not significant. Error bars indicate s.d. (D-E) Snapshots from live cell imaging of WT and S772A/S784A one-cell embryos expressing GFP::H2B and GFP::γ-tubulin. Images show selected stages of cell division in mitosis (D) or meiosis (E). Percentages of embryos displaying a bridging phenotype in the first mitotic division are indicated. Scale bars: 10 μm (D) or 5 μm (E).

We tested whether T750, S772 and S784 can be phosphorylated by CDK-1 *in vitro*. We purified recombinant C-terminal KNL-2 fragments expressed in *Escherichia coli*, with or without substitutions of these three residues to alanines. We then used these recombinant proteins as substrates in an *in vitro* kinase assay (Fig. 1B). The wild type C-terminal KNL-2 fragment was phosphorylated by CDK-1, whereas mutation of all three putative phosphosites to alanines completely prevented phosphorylation. For all single and double S/T to A substitutions, phosphorylation remained detectable. CDK-1 is therefore able to phosphorylate each of the three residues, and does not target other sites within the KNL-2 C-terminal fragment.

Next, to test the role of KNL-2 phosphorylation *in vivo*, we engineered *knl-2* alleles containing substitutions of T750, S772 and S784 to alanines in different combinations using the CRISPR/Cas9 system. Since the deletion of the *knl-2* gene or the depletion of *knl-2* by RNAi leads to severe impairment of cell division and death at early embryonic stages (Maddox et al., 2007), we first scored embryonic lethality levels of the KNL-2 phosphodeficient mutants. We carried out the analysis at two different temperatures: 20°C (permissive temperature), optimal for worm culturing, and 25°C (restrictive temperature), inducing mild temperature stress. The worms bearing single mutation alleles (KNL-2 T750A, S772A or S784A) were superficially wild type, and exhibited wild type levels of embryonic lethality at both tested temperatures (Fig. 1C). However, the double mutant S772A/S784A showed around 25% embryonic lethality at 20°C, and around 75% at 25°C. The triple mutated allele (KNL-2 T750A/S772A/S784A) caused 100% embryonic lethality at both tested temperatures. The strain can be maintained at 15°C, where a few progeny survive to adulthood. The observed increase in severity of the phenotype when combining mutations suggests that these three phosphosites act in a coordinative manner. Since complete loss of KNL-2 function causes fully penetrant embryonic lethality, the KNL-2 T750A/S772A/S784A and S772A/S784A mutants exhibit a partial loss-of-function in addition to being thermosensitive.

To examine the molecular defects underlying the observed embryonic lethality, we focused our further analyses on the strain bearing the KNL-2 S772A and S778A mutations, since it offered the advantage of studying the phenotypic defects at permissive and restrictive temperatures. Using live cell imaging, we examined the progression of the first embryonic division in the S772A/S784A strain expressing GFP::H2B to mark chromatin and GFP::γ-tubulin to mark the spindle poles (Fig. 1D, Movies 1, 2). Compared to the control, the division in S772A/S784A embryos looked defective at several stages: before the nuclear envelope breakdown (NEB), the chromatin appeared more diffuse, indicating potential alterations in chromosome condensation. Cells also exhibited problems in chromosome congression and metaphase plate formation, resulting in disordered metaphase plates. Clear anaphase bridges appeared in all tested cells, indicating errors in chromosome segregation. These defects were apparent at both permissive and restrictive temperatures (Fig. 1D; Fig. S1C), but more pronounced at 25°C (best visible in Movies 1 and 2). Meiotic divisions in the S772A/S784A strain also exhibited anaphase bridges (Fig. 1E). Compromised fidelity of chromosome segregation during meiosis frequently leads to aneuploidy, which is concomitant with the elevated embryonic lethality observed in the S772A/S784A strain. The triple mutant T750A/S772A/S784A exhibited chromosome segregation defects in mitosis and meiosis that are similar to those observed in the S772A/S784A strain even when grown at the permissive temperature for this strain (15°C; Fig. S1D), supporting the hypothesis that all three phosphorylation sites are involved in regulating the same process.

We conclude that KNL-2 residues T750, S772 and S784 are targets for phosphorylation, likely mediated by CDK-1, and work in coordination to regulate KNL-2 functions during cell division.

### KNL-2 functions in centromere and kinetochore formation are not impaired in the S772A/S784A mutant

In *C. elegans* embryos, KNL-2 and CENP-A are interdependent for centromeric localisation and required for kinetochore assembly (Maddox et al., 2007; Oegema et al., 2001). Depletion of *knl-2* by RNAi leads to loss of CENP-A from chromatin and a kinetochore null phenotype, characterised by a complete failure in recruiting other kinetochore proteins (Maddox et al., 2007). We tested if the severe chromosome segregation defects in the S772A/S784A strain resulted from disrupting the KNL-2 function in CENP-A loading or kinetochore formation. We performed these analyses on worms after shifting them to 25°C, where we observed more penetrant phenotypes. We first checked the localisation of KNL-2 and CENP-A (Fig. 2A). In the wild type strain, both proteins were chromatin-bound and followed a similar localisation pattern throughout the division, forming a railroad track-like appearance characteristic for *C. elegans* holocentromeres on prometaphase chromosomes and poleward bi-orientation at metaphase (Buchwitz et al., 1999; Maddox et al., 2007). In the S772A/S784A mutant, these patterns were overall preserved, suggesting that KNL-2 and CENP-A localisation were not defective, although the morphology of the chromosomes appeared altered. In prometaphase, both KNL-2 and CENP-A were chromatin-bound and localised on the face of the chromosomes. The poleward appearance of KNL-2 and CENP-A on metaphase plates was slightly perturbed in the S772A/S784A strain due to some chromosomes not achieving bi-orientation. This observation suggests uncorrected merotelic spindle attachments and is consistent with chromosome bridges detected in the S772A/S784A mutant in anaphase (Fig. 2A).

**Figure 2.**
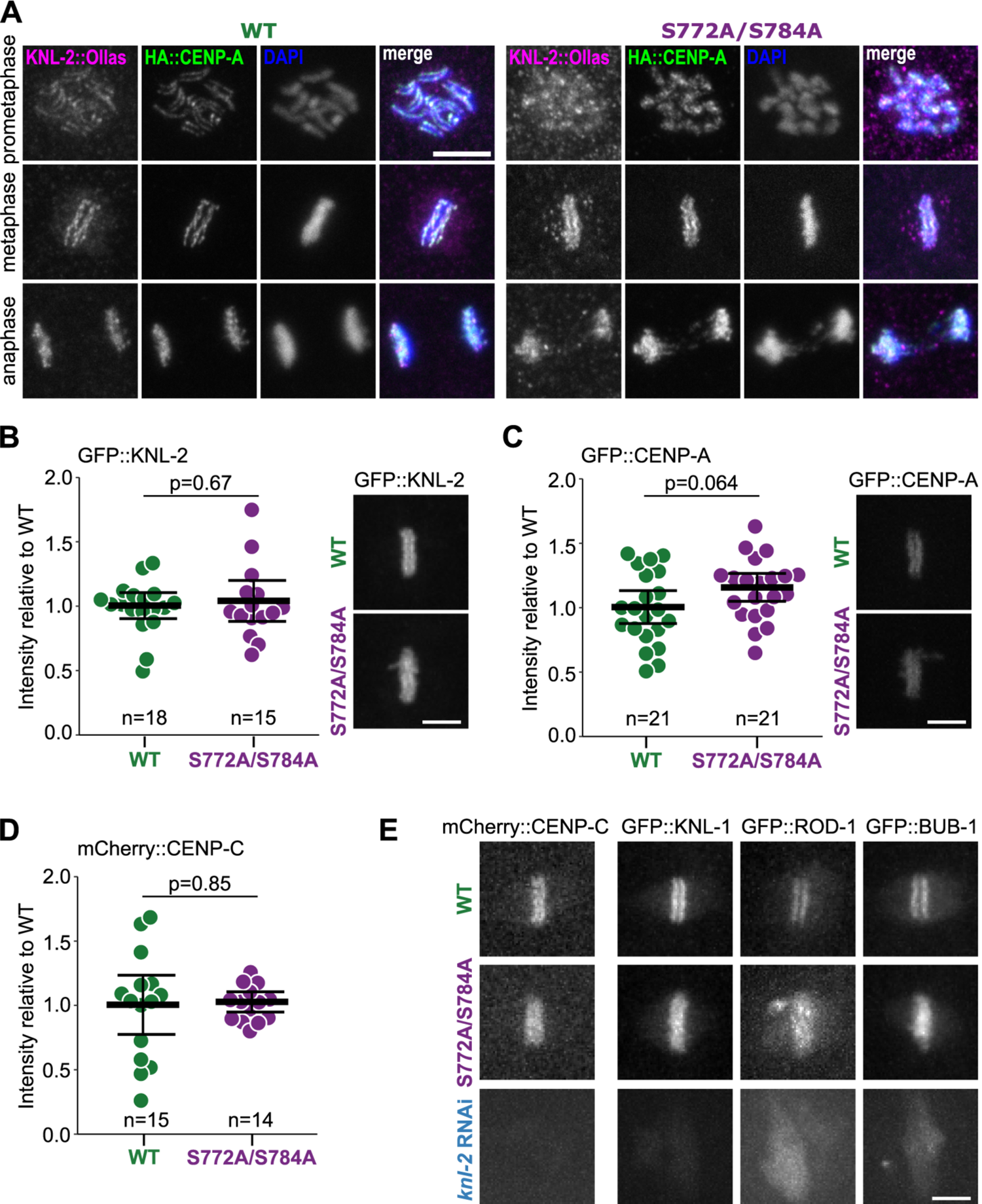
S772A and S784A mutations do not affect KNL-2 localisation, stability, and functions in centromere maintenance and kinetochore recruitment. (A) Immunofluorescent staining of young wild type (WT) and S772A/S784A embryos expressing KNL-2::Ollas and HA::CENP-A at different stages of mitosis. (B-C) Quantification of GFP::KNL-2 (B) or GFP::CENP-A (C) signal on first embryonic metaphases of WT and S772A/S784A strains. Each data point represents one scored embryo. Representative images are shown on the right. (D) Quantification of mCherry::CENP-C signal on first embryonic metaphases of WT and S772A/S784A strains. Each data point represents one scored embryo. Statistical significance was assessed with a t-test (B-D). (E) Representative snapshots from live cell imaging of embryos expressing the mCherry::CENP-C and GFP-tagged kinetochore proteins KNL-1, ROD-1 and BUB-1. Scale bar in all images: 5 μm. In all graphs bars represent mean ± 95% c.i.

Levels of chromatin-bound KNL-2, determined by measuring GFP::KNL-2 signal on mitotic chromosomes at metaphase, were similar in wild type and in S772A/S784A cells, indicating that the phosphosite mutations did not affect protein stability or chromatin association (Fig. 2B). Given that KNL-2 is required for CENP-A loading onto chromatin (Maddox et al., 2007), we measured the metaphase levels of GFP::CENP-A in the S772A/S784A background and found no significant change in comparison to the levels in the wild type strain (Fig. 2C). Phosphorylation of serines 772 and 784 is therefore not involved in regulating the KNL-2 function in CENP-A loading.

Depletion of KNL-2 results in a complete failure to recruit other kinetochore proteins (Maddox et al., 2007). To test if the observed mitotic defects in the S772A/S784A strain are caused by defects in kinetochore assembly, we analysed the localisation of fluorescently tagged kinetochore subunits CENP-C, ROD-1, KNL-1 and BUB-1. CENP-C is the only identified inner kinetochore protein in *C. elegans* and is necessary for the recruitment of all outer kinetochore proteins (Cheeseman et al., 2004; Oegema et al., 2001). mCherry::CENP-C levels on mitotic chromosomes at metaphase were comparable in wild type and S772A/S784A strains (Fig. 2D). CENP-C localisation remained unaffected as well (Fig. S2). The outer kinetochore subunits BUB-1, ROD-1 and KNL-1 also localised to metaphase plates in an expected manner in the S772A/S784A strain (Fig. 2E and Fig. S2), in contrast to almost complete depletion from chromatin after *knl-2* RNAi (Fig. 2E). Although the chromosome alignment defects at metaphase and chromosome bridges suggest kinetochore-spindle attachment defects, kinetochore assembly itself appears normal in the S772A/S784A mutant. We conclude that phosphorylation of KNL-2 on S772 and S784 is not required for centromere formation or kinetochore assembly in *C. elegans* embryos.

### Chromosome condensation is impaired in the S772A/S784A mutant

Previous studies have shown that chromosome condensation is impaired in *C. elegans* cells depleted of KNL-2 or CENP-A (Chan et al., 2004; Hagstrom et al., 2002; Maddox et al., 2007, 2006), but did not address if these condensation defects were a consequence of centromere loss. Since we found that phosphodeficient KNL-2 did not alter CENP-A loading and centromere formation, we considered that it could affect chromosome condensation directly. We therefore investigated the dynamics of chromosome condensation in the S772A/S784A strain in more detail. The changes of GFP::H2B signal distribution in time in the male pronucleus have previously been used as an indicator of the progression of mitotic chromosome formation (Maddox et al., 2006). This method results in a quantifiable condensation parameter that corresponds to the level of chromatin compaction. We followed the mitotic chromosome formation in wild type and S772A/S784A strains at 25°C. Depletion of HCP-6, which encodes a subunit of condensin II complex (Csankovszki et al., 2009; Stear and Roth, 2002), served as a control for chromosome condensation failure (Fig. 3A, B). In the wild type, chromatin condensed steadily, as shown by an almost linear increase of the condensation parameter (Fig. 3A, B). Mitotic chromosomes started to form around 240 s before NEB. In the S772A/S784A mutant, condensation was delayed. An increase in the condensation parameter and visible mitotic chromosome formation appeared only around 90 s before NEB (Fig. 3A, B). We observed a similar delay upon condensin II depletion (*hcp-6* RNAi; Fig. 3A, B).

**Figure 3.**
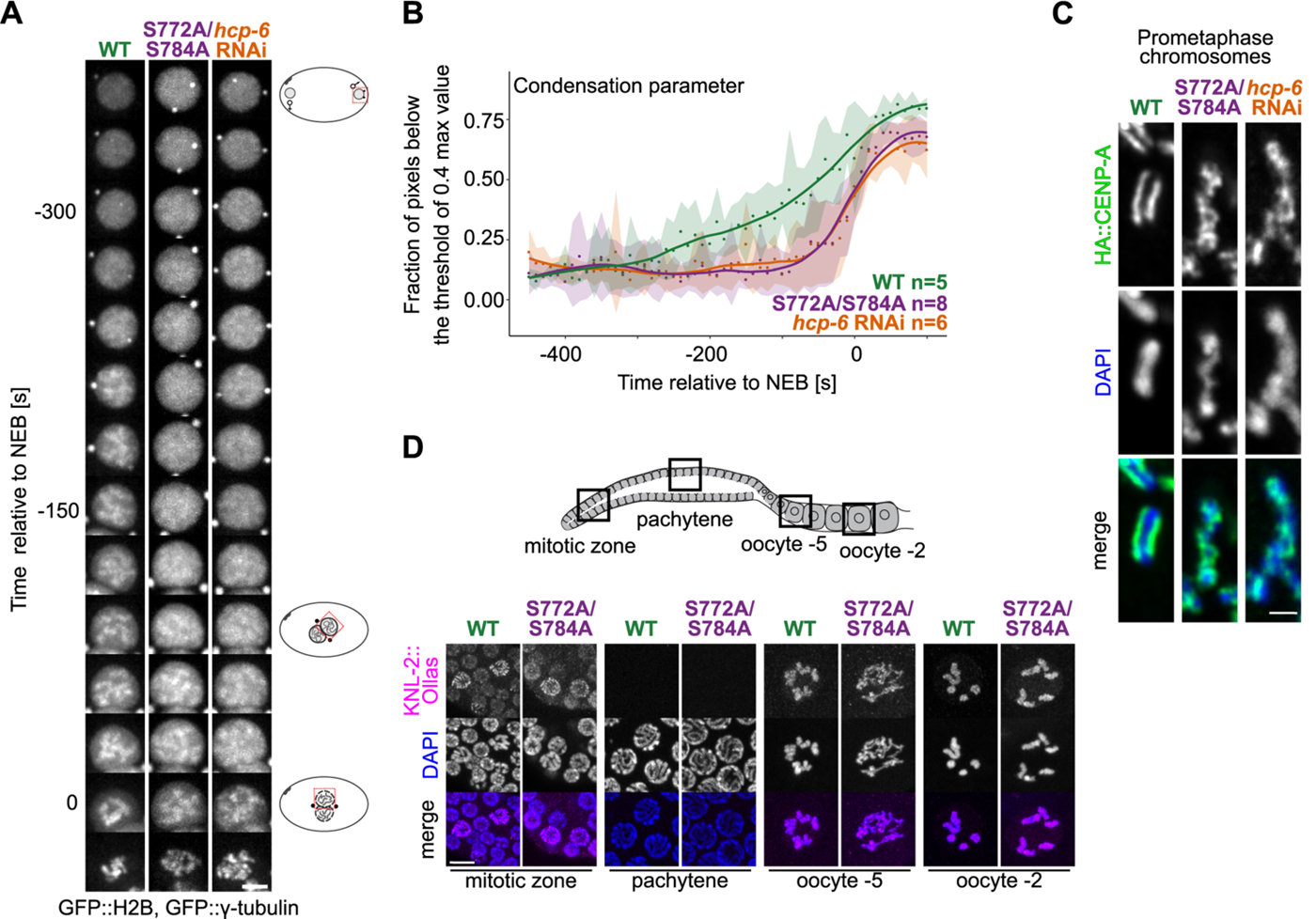
Chromosome condensation is impaired in the S772A/S784A mutant. (A) Kymographs of male pronuclei in embryos expressing GFP::H2B and GFP::γ-tubulin, illustrating the progression of chromosome condensation over time. NEB - nuclear envelope breakdown. Wild type (WT), S772A/S784A, and *hcp-6* RNAi-depletion strains were imaged. Scale bar: 5 μm. (B) Quantification of the condensation parameter in time, for strains as in (A). The condensation parameter is defined as the fraction of pixels with values below an arbitrarily chosen threshold of 0.4 of the maximum pixel value for each image (i. e. timepoint). Dots indicate the average value of the condensation parameter for each timepoint, s.d. is represented as a shaded area. To better illustrate the trend of the condensation parameter changes in time, line plots were fitted with R loess function (span=0.4). n corresponds to the number of embryos scored per condition. (C) Immunofluorescent staining of HA::CENP-A on mitotic prometaphase chromosomes, counterstained with DAPI, in strains as in (A). Each image shows a single chromosome from a prometaphase embryonic cell. Pictures were deconvoluted with the Leica Lightning procedure. Scale bar: 1 μm. (D) *C. elegans* germ line scheme (top), with boxes indicating the location of the nuclei imaged. Immunofluorescence of KNL-2::Ollas in WT and S772A/S784A germ cells, counterstained with DAPI (bottom). Scale bar: 5 μm.

Despite condensation being clearly impaired in the S772A/S784A strain, six separate mitotic chromosomes eventually formed. Their morphology was, however, largely different than in the wild type (Fig. 3A,C). In wild type embryos, chromosomes had a rod-like appearance and looked rigid (Fig. 3C), whereas in the S772A/S784A strain, they appeared more ribbon-like and flexible (Movie 2). Depletion of *hcp-6* by RNAi had a similar effect on the morphology of the chromosomes as the S772A/S784A mutation (Fig. 3C, Movie 3). Close examination of prometaphase chromosomes in fixed samples showed that they were prone to twisting around their own axis in the S772A/S784A strain, resulting in a criss-cross pattern of CENP-A staining, markedly different from the normal railroad track-like CENP-A appearance in the wild type. We observed a similar twisting of chromosomes upon *hcp-6* RNAi, in agreement with previous reports (Stear and Roth, 2002) (Fig. 3C).

The compromised chromosome segregation and increased embryonic lethality in the S772A/S784A strain were already observable at the permissive temperature, albeit at lower levels compared to the restrictive temperature (Fig. 1, Movies 1, 2). To test if the condensation defects also explain the phenotypes at the permissive temperature, we repeated the analysis of the dynamics of chromosome condensation in this condition (Fig. S3 A, B). We found that it was impaired, but seemingly to a lesser degree than at the restrictive temperature (Fig. 3A-B). The condensation defects and consequently the segregation problems are therefore exacerbated rather than triggered by the elevated temperature.

Condensin depletion does not only affect mitosis, but also leads to changes in meiotic chromosome morphology and defects in meiotic divisions (Chan et al., 2004; Houlard et al., 2015; Yu and Koshland, 2003). *C. elegans* germ cells move through the gonad from distal to proximal zone as they progress through meiosis. CENP-A and KNL-2 are present on chromatin in the mitotically proliferating zone (distal zone), then they are removed at the onset of meiosis (transition zone) and associate with chromatin again in diplotene (proximal zone) (Gassmann et al., 2012; Maddox et al., 2007; Prosée et al., 2020). We investigated chromosome morphology in the S772A/S784A meiotic germ cells and found that it was altered in the proximal zone of the germ line, but seemed unaffected in the pachytene zone (Fig. 3D). The observed meiotic condensation defects were thus only observed in regions of the germ line where KNL-2 and CENP-A are present. Early diakinetic nuclei typically contain well defined bivalents, but in the S772A/S784A strain individual chromosomes were hardly distinguishable at these stages (Fig. 3D). The differences between the S772A/S784A mutant and the wild type became less pronounced as oocyte maturation progressed, and in late diakinetic oocytes, six individual chromosome tetrads were visible in both strains (Fig. 3D). The localisation of KNL-2 remained unaffected, similarly to what we observed for embryonic cells (Fig. 3D). Our results suggest that phosphorylation of the KNL-2 C-terminus is not only regulating mitotic chromosome condensation, but plays a similar role in meiotic diplotene and diakinesis stages.

Condensation failure has previously been shown to cause segregation problems (Chan et al., 2004; Csankovszki et al., 2009; Hagstrom et al., 2002; Hudson et al., 2003; Maddox et al., 2006; Oliveira et al., 2005; Ono et al., 2004; Samoshkin et al., 2009; Steffensen et al., 2001). To assess whether the aberrant segregation in the S772A/S784A strain is a consequence of faulty mitotic chromosome formation, we examined the first embryonic mitoses in this strain and after *hcp-6* RNAi in the wild type strain (Fig. S3C-E). To compare the progression of the division, we examined the NEB-to-anaphase onset interval and measured the spindle pole distance over time (Fig. S3C,D). Both S772A/S784A mutation and depletion of *hcp-6* caused similar defects: chromosome congression problems and clear anaphase bridges (Fig. S3E, Movies 2, 3). The kinetics of spindle pole separation was comparable in *hcp-6* depleted embryos and in the S772A/S784A strain, showing slightly premature pole separation relative to wild type (Fig. S3D). This premature pole separation may be a consequence of delayed formation of attachments between kinetochores and microtubules, rather than of defects in kinetochore assembly, as we found that kinetochore proteins localise normally in the S772A/S784A strain (Fig. 2D, E). The time between NEB and anaphase onset was comparable for both S772A/S784A strain and *hcp-6* depletion, and longer than in the wild type (on average 198 s for S772A/S784A, 212 s for *hcp-6* RNAi, and 151 s in the wild type), as expected from the observed defects in chromosome congression and formation of the metaphase plate (Fig. S3 E). Together, these results indicate that depletion of *hcp-6* causes mitotic defects reminiscent of the ones observed in the S772A/S784A strain, suggesting that the segregation defects are indeed a downstream consequence of condensation problems. We conclude that phosphorylation of KNL-2 S772 and S784 is required for chromosome condensation during meiosis and at the onset of mitosis.

### Condensin II levels are reduced on metaphase chromosomes in the S772A/S784A mutant

Since we found that the combined mutation of KNL-2 S772 and S784 to alanines affected chromosome condensation independently of centromere formation, and that depletion of condensin II resulted in similar phenotypic defects, we hypothesised that phosphorylation of S772 and S784 might be required for condensin recruitment or maintenance. As in most eukaryotes, the *C. elegans* genome encodes two condensin complexes responsible for mitotic chromosome formation, called condensin I and condensin II (Csankovszki et al., 2009). Each complex consists of five subunits: two structural maintenance of chromosomes (SMC) subunits and three accessory subunits (Csankovszki et al., 2009; Ono et al., 2003). While SMC subunits are shared by both condensin complexes, the accessory subunits are unique for each condensin complex: DPY-28 (CAP-D2), CAPG-1 (CAP-G) and DPY-26 (CAP-H) for condensin I, and HCP-6 (CAP-D3), CAPG-2 (CAP-G2) and KLE-2 (CAP-H2) for condensin II (Csankovszki et al., 2009; Hirano, 2012). We used GFP-tagged KLE-2 for visualising condensin II and GFP-tagged CAPG-1 for condensin I. Both complexes exhibited the same subcellular localisation and nuclear dynamics in wild type and S772A/S784A embryos (Fig. 4A, Movies 4, 5). Condensin II localised to the nucleus already in prophase, whereas condensin I associated with the forming chromosomes after NEB. Both complexes remained chromatin bound for the remaining stages of mitosis. The observed mitotic defects in the S772A/S784A mutant are therefore unlikely to be caused by altered timing of condensin localisation.

**Figure 4.**
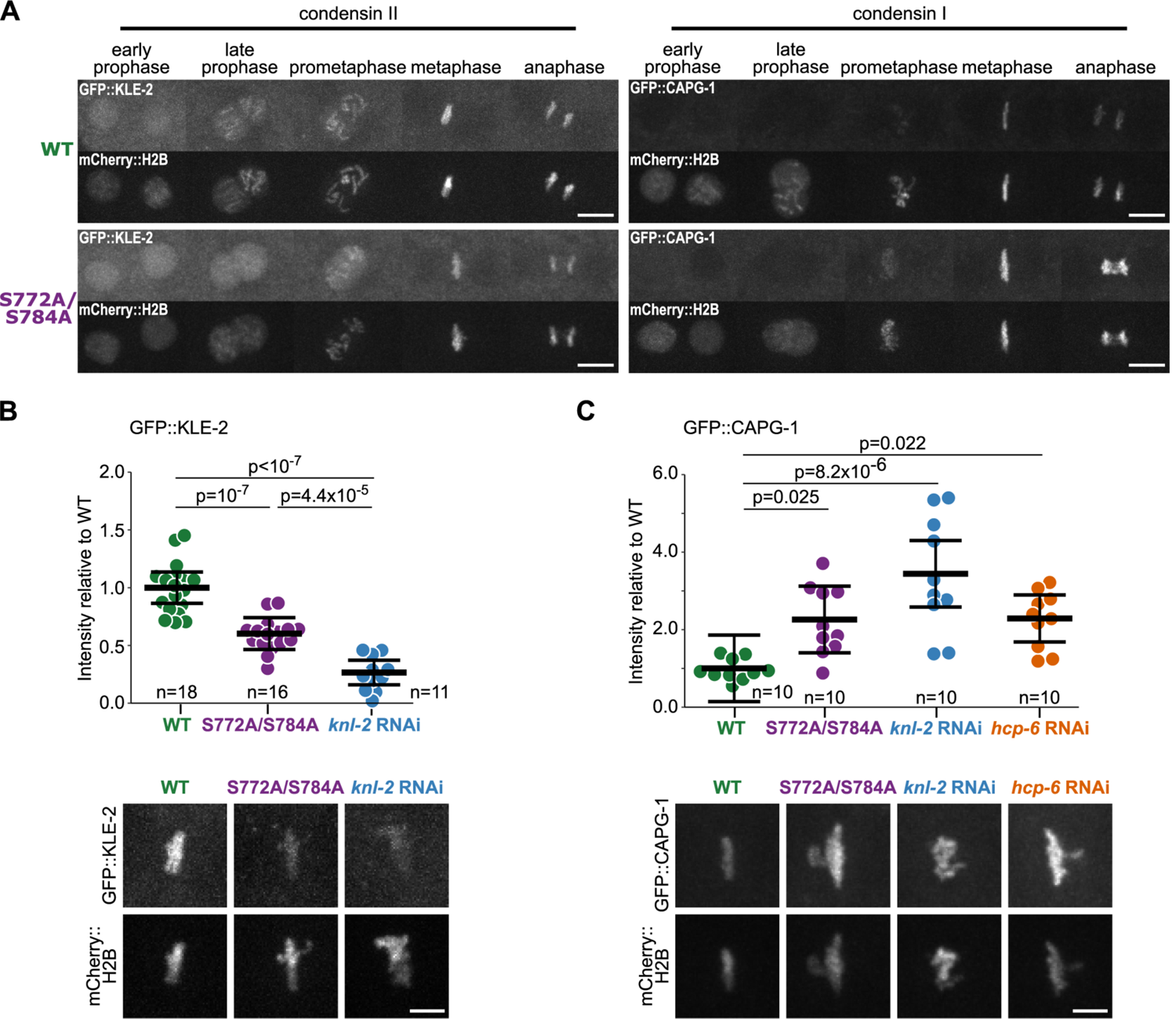
Condensin complex levels on chromatin are changed in the S772A/S784A mutant. (A) Images of selected stages of first embryonic mitosis in wild type (WT) and S772A/S784A strains expressing GFP::KLE-2 (condensin complex II) or GFP::CAPG-1 (condensin complex I) and mCherry::H2B. Scale bar: 10 μm (B-C) Quantification of GFP::KLE-2 (B) and GFP::CAPG-1 (C) signal on first embryonic metaphases, in indicated strains. Each data point represents one scored embryo. Representative images are shown below the quantifications. Scale bar: 5 μm. For assessing statistical significance ANOVA followed by the Tukey-Kramer test was used. In all graphs bars represent mean ± 95% c.i.

We next tested if S772A/S784A mutants were defective in condensin recruitment or maintenance. We quantified the levels of GFP::KLE-2 and GFP::CAPG-1 on mitotic chromosomes at metaphase. The mean levels of GFP::KLE-2 were significantly lower in the S772A/S784A mutant compared to the wild type (around 60% of the wild type level) (Fig. 4B). This result suggests that phosphorylation of KNL-2 is involved in regulating condensin II levels on mitotic chromosomes. Consistent with this observation, depletion of *knl-2* by RNAi also resulted in reduced levels of GFP::KLE-2, to an even greater degree than S772A and S784A mutations (26% of the wild type level). Total GFP::KLE-2 levels in embryonic lysates were similar in the wild type and in the S772A/S784A strain, as determined by western blot (Fig. S4A), indicating that the stability of the condensin II complex was not affected. Over the course of interphase, the nuclear levels of GFP::KLE-2 were comparable in both strains (Fig. S4B), suggesting that KLE-2 import into the nucleus was not majorly affected either. We therefore conclude that a specific phosphorylation status of the KNL-2 C-terminal region is required for condensin II to properly associate with chromatin during mitotic chromosome formation. Similar effects of the S772A/S784A mutations on GFP::KLE-2 levels were visible at the permissive temperature (20°C, Fig. S4C), concomitant with the condensation defect observed in this condition.

We next tested if the S772A/S784A mutations also affected the condensin I complex. Condensin I levels, measured by GFP::CAPG-1 signal on metaphase plates, were elevated in the S772A/S784A embryos in comparison to the wild type (Fig. 4C), while total protein levels in embryonic lysates remained unchanged (Fig. S4A). Depletion of *knl-2* by RNAi also resulted in increased GFP::CAPG-1 levels at metaphase. This increase is likely a downstream consequence of decreased condensin II levels, since depletion of *hcp-6* by RNAi mirrored this effect (Fig. 4C). These observations suggest that condensin I levels increase to compensate for the insufficient levels of condensin II on mitotic chromosomes.

We conclude that lack of phosphorylation of KNL-2 S772 and S784 leads to reduced levels of condensin II on mitotic chromosomes. This reduction of condensin II explains the condensation defects and altered chromosome morphology observed in S772A/S784A mutant embryos and the proximal part of the germ line, and likely underlies the chromosome segregation defects and the embryonic lethality.

## DISCUSSION

### The role of *C. elegans* KNL-2 in chromosome condensation is distinct from its function in CENP-A loading

We found that the centromeric protein KNL-2 is involved in regulating chromosome condensation in the holocentric nematode *C. elegans* through the phosphorylation status of its C-terminal region established by CDK-1. Abolishing this phosphorylation results in reduced levels of condensin II on mitotic chromatin, aberrant chromosome morphology and segregation defects, but has no effect on CENP-A loading and kinetochore assembly. Our results suggest that KNL-2 has taken on a direct role in regulating mitotic chromosome formation, and that this role is independent from its function in centromere maintenance.

CDK-1-dependent phosphorylation also regulates vertebrate homologues of KNL-2, M18BP1, in human cells (McKinley and Cheeseman, 2014; Pan et al., 2017; Silva et al., 2012; Spiller et al., 2017; Stankovic et al., 2017), and *Xenopus laevis* (French and Straight, 2019). M18BP1 is a subunit of the MIS18 complex required for the initiation of CENP-A loading (French and Straight, 2017; Fujita et al., 2007; Hayashi et al., 2004; Hori et al., 2017; Moree et al., 2011; Pan et al., 2019). Phosphorylation of human M18BP1 by CDK-1 regulates MIS18 complex formation and localisation ensuring the proper timing of CENP-A loading (Foltz et al., 2009; Pan et al., 2017; Spiller et al., 2017).

Although KNL-2 is also indispensable for CENP-A loading in worms (Maddox et al., 2007), we found that in contrast to the role of M18BP1 phosphorylation in vertebrates, CDK-1-dependent phosphorylation of KNL-2 S772 and S784 is not required for CENP-A loading (Fig. 2), but instead for chromosome condensation (Fig. 3). These KNL-2 residues are located in a C-terminal region that is not conserved between nematodes and vertebrates, but is conserved within the *Caenorhabditis* genus (Fig. S1A), indicating that this phosphoregulation and its role in chromosome condensation might be present also in other holocentric nematodes.

### The regulation of chromosome condensation by centromeres

Indication that the centromeres might be involved in regulating chromosome condensation in *C. elegans* comes from early work where disrupting centromeric chromatin by depleting CENP-A or KNL-2 has been shown to lead to dramatic condensation defects (Chan et al., 2004; Hagstrom et al., 2002; Maddox et al., 2007, 2006). Aside from *C. elegans*, a regulatory role of centromeres in mitotic chromosome condensation has only recently been described in *Saccharomyces cerevisiae*, where centromeres act as the initiation sites for condensation, triggering a regulatory cascade that spreads along the chromosome arms (Kruitwagen et al., 2018). For holocentric chromosomes, such a spreading mechanism would appear redundant, as centromeres are distributed along the length of the entire chromosomes.

In agreement with previous observations, we show that the centromeric protein KNL-2 is involved in chromosome condensation, as it regulates condensin II levels on chromatin. However, we found that CENP-A loading is preserved in the S772A/S784A strain (Fig. 2), suggesting that correctly maintained centromeric chromatin is insufficient to ensure proper condensin II targeting. The notion that chromosome condensation is regulated independently of CENP-A in *C. elegans* is supported by the observations that the depletion of either of the SMC subunits SMC-4 and MIX-1 does not prevent CENP-A chromatin binding (Chan et al., 2004; Hagstrom et al., 2002) and depletion of CAPG-1 or CAPG-2 (condensin I or II, respectively) has no detectable effect on CENP-A levels on endogenous chromosomes (Lin and Yuen, 2021). It is likely that the previously observed condensation defects upon CENP-A depletion were caused by the simultaneous loss of KNL-2, the C-terminal phosphorylation status of which is crucial to regulate the mitotic chromosome formation.

### KNL-2 C-terminal phosphorylation regulates condensin complex levels on mitotic chromosomes

The reduced levels of the condensin II complex on chromosomes in the phosphodeficient KNL-2 strain (Fig. 4) could be caused by delayed or less efficient condensin II recruitment to chromatin. Alternatively, the condensin II complex might be properly recruited, but not retained on chromatin in S772A/S784A mutants due to weaker or less stable chromatin binding. Considering that condensin II achieves a stable chromatin association during prophase in human cells (Gerlich et al., 2006), we favour the hypothesis of condensin II being less efficiently recruited to chromatin in the S772A/S784A strain. It has recently been observed that when KNL-2 loses its interaction with CENP-A, it fails to localise to chromatin, and chromosome condensation becomes defective (Prosée et al., 2020). This suggests that KNL-2 chromatin association precedes condensin II recruitment.

Condensin I targets chromatin independently from condensin II (Green et al., 2012; Hirota et al., 2004; Ono et al., 2003) and performs distinct functions in mitotic chromosome formation (Gibcus et al., 2018; Ono et al., 2004). The elevated levels of condensin I on metaphase chromosomes in the S772A/S784A strain are likely a secondary effect of the condensin II depletion from chromatin, as we observed a similar effect after *hcp-6* RNAi (Fig. 4). Condensin I over-recruitment may compensate for partial depletion of condensin II. It is possible that when one of the complexes is depleted, some condensin binding sites on chromatin are left unoccupied, and the other complex binds or spreads to them instead. A similar effect has been observed in *X. laevis*, where after condensin I depletion, condensin II became enriched on chromosome arms (Hirota et al., 2004). We note, however, that the condensin I marker used here (CAPG-1) has other documented roles during cell division, as it accumulates on chromosomal bridges, spindle midzone and midbody in later stages of mitosis (Bembenek et al., 2013). We therefore cannot exclude that the increased presence of CAPG-1 on the metaphase chromosomes is in relation to its other functions, rather than to its canonical condensin I roles.

In the T750A/S772A/S784A strain, which lacks an additional phosphorylation site at the KNL-2 C-terminus, the cell division defects are more severe than in the S772A/S784A mutant (Figs. 1, S1). These defects are likely caused by further decrease in condensin II levels and more pronounced condensation impairment. However, in contrast to the fully penetrant embryonic lethality upon condensin II depletion, the T750A/S772A/S784A strain is viable at 15°C, suggesting that condensin II recruitment to the chromatin is not completely abolished when the KNL-2 C-terminus is not phosphorylated. Therefore, additional unidentified KNL-2 phosphosites might be involved in the regulation of chromosome condensation. Alternatively, KNL-2-dependent condensin II recruitment could be redundant with other regulatory pathways.

### Chromosome morphology is sensitive to condensin II levels and correlates with downstream defects

Due to decreased levels of condensin II complex on chromatin, mitotic chromosomes in the S772A/S784A strain lacked normal rigidity and were prone to twisting around their axis (Figs. 3, 4). In other species the depletion of condensin complexes also did not lead to total condensation failure, but rather affected the physical properties of chromosomes (Gerlich et al., 2006; Hagstrom et al., 2002; Hudson et al., 2003; Oliveira et al., 2005; Ono et al., 2003; Steffensen et al., 2001). Specifically, condensin II depletion resulted in elongation of the chromosomes in human (Gibcus et al., 2018; Ono et al., 2017, 2003) and chicken cells (Green et al., 2012). In *C. elegans*, HCP-6 depletion caused chromosomes to twist around their axis (Stear and Roth, 2002), a result reproduced in this study (Fig. 3). Given that we found no evidence for centromere defects in the S772A/S784A strain, we attribute all its cell division defects to this impaired chromosome morphology. We observed problems with chromosome congression, bi-orientation, and anaphase bridges, which resemble the defects observed upon *hcp-6* depletion (Fig. S3). Consistent with our observations, condensin depletion frequently leads to defects in metaphase plate formation and anaphase bridging (Chan et al., 2004; Csankovszki et al., 2009; Hagstrom et al., 2002; Hudson et al., 2003; Maddox et al., 2006; Oliveira et al., 2005; Ono et al., 2004; Samoshkin et al., 2009; Steffensen et al., 2001).

### KNL-2 exhibits a similar role in meiotic and mitotic chromosome condensation

Chromosome condensation is essential not only for mitotic, but also for meiotic divisions. We observed segregation defects in meiosis (Fig. 1) and impaired chromosome morphology in the proximal zone of the germ line (Fig. 3) in the S772/S784A strain. Interestingly, both KNL-2 and condensin II subunits show a discontinued pattern of chromatin association in the *C*. *elegans* germ line. KNL-2 is present in the mitotic zone, then removed at the entry to meiotic prophase and reappears when the cells reach the diplotene stage of meiotic prophase (Prosée et al., 2020). The condensin II complex, although present in all germline nuclei, is chromatin-bound only in the mitotic zone, diplotene and diakinesis (Chan et al., 2004). The apparent overlap between the stage when KNL-2 and condensin II re-associate with chromatin and the onset of condensation defects in the S772A/S784A strain suggests that the same mechanism of KNL-2-dependent condensin II recruitment is compromised in S772A/S784A strain in mitosis and meiosis. Consistent with our observations, HCP-6 mutants exhibit condensation defects in diplotene-stage nuclei, but not in pachytene nuclei (Chan et al., 2004). A recent study found that CDK-1 acute depletion causes chromosome morphology changes in the proximal zone of the germ line (Brandt et al., 2020). These changes are similar to those observed in the S772A/S784A strain, further strengthening our hypothesis of CDK-1 involvement in the regulation of KNL-2 function in chromosome condensation.

Taken together, we show that the centromeric protein KNL-2 plays a direct regulatory role in chromosome condensation. We propose that KNL-2 integrates the crosstalk between the condensation machinery and the centromeres. The separation-of-function alleles generated in this study allow to disentangle the roles of KNL-2 in centromere maintenance and chromosome condensation, and are a first step towards deciphering the mechanism of centromere involvement in chromosome condensation in *C. elegans*.

## MATERIALS AND METHODS

### Worm maintenance

Worms were cultured according to standard procedures (Brenner, 1974) on NGM (nematode growth medium) plates seeded with OP50 bacteria The worms were maintained at 20°C or 15°C (for ts strains) and shifted to 25°C for analysis, as indicated.

### Strain construction

Strains are listed in Table S2. CRISPR/Cas9 was used for modifying endogenous loci as described previously (Arribere et al., 2014). For the point mutation T750A and small tag insertion (OLLAS, HA), oligonucleotides were used as repair templates. For mutating serines 772 and/or 784, two cuts were introduced, and a *knl-2* fragment (of 712 bp) containing the mutations and a C-terminal OLLAS tag was PCR-amplified from pJW56 or pJW57 plasmids (Table S3) and used as a repair template. For introducing fluorescent tags (GFP and mCherry) the repair templates were generated by PCR with primers containing ∼50bp overhangs with homology to the target locus. Transgenic *gfp::knl-2* constructs were introduced by MosSCI (Frøkjær-Jensen et al., 2014). Genomic sequence of *knl-2* locus with S772A and S784A mutations, its promoter and 3’UTR regions were cloned into the pCFJ151 vector together with 3xFLAG and GFP coding sequences. Table S3 contains the names of plasmids and sequences of sgRNAs used for strain generation. All other strains were obtained by genetic crossing.

### Embryonic viability assessment

10 L4 hermaphrodites were singled on NGM plates seeded with a small amount of OP50 bacteria and maintained at 20°C or 25°C until the young adult worms laid 20-40 eggs. The worms were then removed and the number of eggs counted. The plates were incubated for another 20-30 h and the L1 hatchlings were counted. The embryonic lethality was assessed as the difference between the number of laid eggs and the number of hatchlings. The experiment was repeated three times for each condition.

### RNA interference

Bacteria expressing specific dsRNAs were obtained from the Ahringer library (Source BioScience) and the RNAi experiments were performed by feeding as described previously (Kamath et al., 2001) with minor modifications. Briefly, an overnight culture of dsRNA expressing bacteria was diluted 50 times (to OD∼0.05-0.1) in LB media containing ampicillin. Expression of dsRNA was induced with 1 mM IPTG when the culture reached OD∼0.6-0.8. After 4 h, bacteria were concentrated 50 times and used to seed NGM plates supplemented with 1 mM IPTG and carbenicillin. L4 hermaphrodites were put on RNAi plates for 18-20 h at 25°C or 24 h at 20°C.

### Antibodies and western blotting

For generating embryonic lysates, gravid adult hermaphrodites were bleached to release embryos. The embryo pellet was resuspeded in 2-3 volumes of lysis buffer (8 mM Na_2_HPO_4_, and 2 mM KH_2_PO_4_ 137 mM NaCl, 100 mM KCl, 1mM MgCl_2_, 1 mM EGTA, 10% glycerol, 1% CHAPS, 1 mM PMSF) and snap-frozen in liquid nitrogen. To break the embryo shells and shear chromatin, the samples were sonicated with the Bioruptor Pico (Diagenode) machine (10 cycles: 30s ultrasound, 30s rest) with occasional re-freezing in liquid nitrogen. The lysates were cleared by centrifugation (21’000 g, 15 min) and the protein amount was quantified with the use of Bio-Rad Protein Assay (Bio-Rad, 5000006). For western blotting the following primary antibodies were used: anti-tubulin (Abcam, ab6160), anti-GFP (Abcam, ab290). The LI-COR Odyssey system with fluorescent secondary antibodies (IRDye) was used for detection.

### Immunoprecipitation, mass spectrometry and phosphosites identification

Immunoprecipitations were performed as described previously (Prosée et al., 2020). Briefly, embryos obtained by bleaching were snap-frozen in RIPA buffer (50 mM Tris-HCl pH = 7.4, 500 mM NaCl, 0.25% deoxycholate, 10% glycerol, 1% NP-40, 2 mM DTT, EDTA-free protease inhibitor cocktail (Roche), PhosSTOP (Roche)), sonicated for 15 cycles, and lysates cleared by centrifugation for 30 minutes (sonication and centrifugation parameters as above). Lysates were incubated overnight at 4°C with Pierce Anti-HA Magnetic Beads (Thermo Fisher Scientific). Beads were then washed, and boiled in Pierce Lane Marker Non-Reducing Sample Buffer (Thermo Fisher Scientific). The eluates were analysed by the Proteomic Facility at the Functional Genomics Center Zurich. Samples were processed according to standard procedures used by the facility. The proteins were precipitated with trichloroacetic acid, washed with acetone, resuspended and digested with trypsin. Samples were then dried, dissolved in 0.1% formic acid and ca. 10% of the sample was injected into the LC/MS/MS system. The raw files produced by the spectrometer were processed with MaxQuant version 1.6.0.16 (Cox and Mann, 2008). Peptide searches were run against the *C. elegans* proteome (UP000001940) with the following parameters: minimal peptide length: 7 aa, maximum 2 missed cleavages (trypsin/p digestion), False Discovery Rate: 0.05, modifications: N-terminal acetylation, methionine oxidation and phosphorylation (STY), maximum 5 modifications per peptide allowed. Identified Peptide Spectrum Matches containing a putative phosphorylation and mapping to KNL-2 were then further manually inspected.

### Expression and purification of recombinant proteins

The sequence encoding the C-terminal KNL-2 fragment (residues 617-877) was cloned into a vector derived from pET allowing for expression in *E. coli* as a fusion protein with a TEV cleavage site and a His_6_-GST tag at the C-terminus. The mutations T750A, S772A and S784A in different combinations were introduced by the Quick Change Mutagenesis Kit (Stratagene). The recombinant proteins were overexpressed in *E. coli* Rosetta2, grown in 2xYT media at 37°C for 4 h followed by overnight induction at 18°C with 0.1 mM isopropyl-β-D-thiogalactopyranoside (IPTG). Induced cells were harvested by centrifugation and resuspended in lysis buffer (50 mM phosphate buffer pH 8.0, 600 mM NaCl, 10% glycerol, 25 mM imidazole, 0.15% CHAPS, 5 mM β-mercaptoethanol, 1 μg/ml DNase, 1 μg/ml lysozyme, 1 mM PMSF, 1 μg/ml leupeptin, and 2 μg/ml pepstatin). Cells were lysed using an Emulsiflex system (AVESTIN) and cleared by centrifugation at 15 000 rpm for 45 min at 4°C. The soluble fraction was subjected to an affinity purification using a chelating HiTrap FF crude column (GE Healthcare) charged with Ni^2+^ ions on an AKTA-HPLC purifier (GE Healthcare). The proteins were washed with lysis buffer containing 300 mM NaCl, and eluted with 300 mM NaCl and 250 mM imidazole. The purest fractions were combined and passed over a desalting column (GE Healthcare). The TEV cleavage was performed overnight at 8°C with a His_6_-tagged TEV protease at a ratio of 1:20. The sample was reloaded on the Ni-NTA column and the flow-through containing the pure protein was collected. Samples were concentrated (Amicon 30 kDA) and loaded on a Superdex GF75 Increase column. The pure proteins were concentrated to about 1-1.2 mg/ml.

### Kinase Assay

Each reaction (total volume: 10 μl) contained 1 μg recombinant protein, 0.25 mM cold ATP, 5 μCi γ^32^P-ATP and 350 ng CDK1/Cyclin B Recombinant Human Protein (Thermo Fisher PV3292) in kinase buffer (50 mM HEPES pH 7.5, 10 mM MgCl2, 1 mM EGTA, 0.01% Brij-35) supplemented with PhosSTOP (Sigma-Aldrich) and Complete EDTA-free proteases inhibitors (Sigma-Aldrich). The samples were incubated at 30°C for 10 min. The reactions were stopped by adding 3x Laemmli sample buffer and boiling at 95°C for 5 min. The samples were then resolved on a 10% acrylamide gel. The gel was stained with Coomassie, dried and exposed to a phosphoimager screen (GE Healthcare). The results were analysed with a Typhoon FLA 9500 (GE Healthcare).

### Staining and imaging of fixed samples

Young adult hermaphrodites were washed twice in PBS with 0.1% Triton X-100 to remove bacteria, then cut in half in anesthetizing buffer (50 mM sucrose, 75 mM HEPES pH 6.5, 60 mM NaCl, 5 mM KCl, 2 mM MgCl_2_, 10 mM EGTA pH 7.5, 0.1% NaN_3_). The released embryos were transferred onto glass slides coated with poly-L-lysine. A coverslip was placed on top and the samples were freeze-cracked and fixed in cold (−20°C) methanol for 5 min. After two washes in PBS (5 min each) samples were incubated with primary antibodies overnight at 4°C (anti-HA antibody mAb 42F13, 1:60; anti-OLLAS antibody Novus Biologicals NBP1-06713B, 1:150). Slides were washed two times in PBS, incubated with secondary antibody (Alexa Fluor 488 goat anti-mouse, Alexa Fluor 594 goat anti-rat; 1:700) for 1-2 h at room temperature and counterstained with DAPI (2 μg/ml) for 15 min. Samples were washed once in PBS and mounted in VECTASHIELD Antifade Mounting Medium (Vector Laboratories). Images were taken with a Leica SP8 confocal microscope, using a 100x oil objective (aperture: 1.40). Z-stacks with 0.3 μm steps were acquired. For some images the Leica SP8 LIGHTNING function was used for image deconvolution (as indicated in the figure description). Images were processed with Fiji software (Schindelin et al., 2012): contrast adjusted for display, maximum intensity Z-projection, Gaussian blur filter (radius: 0.5 pixel).

### Live imaging of embryos

L4 hermaphrodites were placed on NGM plates or RNAi plates at 25°C for 18-20 h or 24 h at 20°C before imaging. Young adult worms were cut in egg buffer (118 mM NaCl, 48 mM KCl, 2 mM CaCl_2_, 2 mM MgCl_2_ and 25 mM HEPES pH7.5), and released embryos were mounted on 2% agarose pads. The imaging was performed on a spinning disc confocal system (Intelligent Imaging Innovations Marianas SDC) mounted on an inverted Leica DMI microscope (Photometrics Evolve 512) with 63x oil objective (aperture: 1.4). The microscope was equipped with a temperature chamber set to 25°C or 20°C, depending on the experiment. For each time series, 10-12 Z-sections of 0.8 µm were taken every 10 s with 1×1 binning. The Z-position was adjusted manually during the imaging. Lasers were set to 100% power, the camera intensity was set to 800 and the gain to 3. The exposure time was determined for each fluorescent protein separately. The images were analysed with Fiji software with some steps partially automated with custom macros (see below).

### Quantification of fluorescence intensity on metaphase plates in one-cell embryos

Total intensity of fluorescently-tagged proteins was determined for a time series acquired as described above. For strains expressing GFP::KLE-2/mCherry::H2B and GFP::CAPG-1/ mCherry::H2B, images from both channels were recorded simultaneously. Metaphase plate was defined as the last frame before the anaphase onset, which was defined as the first frame when sister chromatids appeared to separate. For segmentation, the maximum intensity Z-projections of the mCherry::H2B images were automatically thresholded using the MaxEntropy function in Fiji. The created mask was used to select a region of interest (ROI) spanning the metaphase plate. Next, this ROI was expanded by 5 pixels in each direction to calculate the background levels. SUMstack projections of the corresponding GFP images (GFP::KLE-2 or GFP::CAPG-1) were used for measurements of total intensity of each defined ROIs. Then, the mean background value was determined for each measurement by subtracting the total intensity of the metaphase ROI from the total intensity of the expanded ROI and dividing the resulting value by the difference in ROI area. The total background for each measurement was calculated as the metaphase ROI area multiplied by the respective mean background value. Total background was then subtracted from the total intensity of the metaphase plate ROI and the resulting values were used in statistical analysis as described below. The analysis of strains expressing only one fluorescent marker (GFP::CENP-A, GFP::KNL-2 or mCherry::CENP-C) was done similarly, but the single channel was used for both defining the ROIs and measuring the intensities.

### Quantification of total GFP::KLE-2 nuclear fluorescence signal in one-cell embryos

Total nuclear intensity of GFP::KLE-2 was determined for time series acquired as described above. Since these conditions do not ensure that the entire volume of both pronuclei is encompassed within each Z-stack, maximum intensity Z-projections were used for estimations rather than SUMstacks projections. For segmentation, mCherry::H2B images were contrast adjusted, mean filter (2 pixel radius) was applied, and images were automatically thresholded using MaxEntropy function in Fiji. Created masks were used to determine ROIs spanning maternal and paternal pronucleus. ROIs were then used to measure the total intensity in the corresponding GFP::KLE-2 image. Background was estimated by manually choosing a region (25×25 pixels) within the cytoplasm and measuring mean pixel value for each image from the time series. This value was then multiplied by total nuclear ROI area and subtracted from the sum of integrated intensities for maternal and paternal pronucleus. Resulting corrected total nuclear intensities were divided by total ROI area to obtain mean pixel values displayed in graphs.

### Quantification of pole to pole distance in one-cell embryo

Strains expressing γ-tubulin::GFP and GFP::H2B were used for determining the spindle poles separation over time. Time series were acquired as described above, maximum intensity Z-projections were used for measurements. The distance between spindle poles was measured manually with a line tool in Fiji. The measurements were aligned in time relative to the anaphase onset defined as the first frame when sister chromatids began to separate.

### Quantification of cell cycle timing

The time between NEB and anaphase onset was measured using the same images as for the pole to pole separation. NEB was visually determined as the moment when the signal intensity of nucleoplasmic GFP::H2B becomes indistinguishable from the cytoplasmic background. Anaphase onset was the first frame when sister chromatids began to separate.

### Quantification of condensation

Quantification of the condensation parameter was performed as described previously (Maddox et al., 2006). Images were segmented as for total nuclear signal estimation, except that the Otsu thresholding function was used. Then, a square ROI of 21×21 pixels was centred on the paternal nucleus. Distribution of pixel values within each ROI was obtained with Fiji histogram function.

## Statistical analysis

The values from intensity measurements were normalised by dividing each value by the average of the control measurements. This sets the average for wild type measurements to 1 in each case. The data were processed and plotted with the use of R plyr, FSA and ggplot2 packages (Ogle et al., 2021; Wickham, 2011, 2009). The statistical analysis was performed according to guidelines in (Pollard et al., 2019) and the custom scripts were based on suggestions therein. Briefly, the normal distribution of values was assessed by plotting histograms of values for each sample and running a Shapiro-Wilk test. Next, the variances were calculated for each sample. If the data were distributed normally (Shapiro-Wilk p-value<0.05), the following tests were used: two-sided Student’s t-test for comparing two samples of similar variances (difference smaller that 3-fold), Welch’s t-test for two samples with different variances, two-sided ANOVA for comparison of more than two samples. ANOVA was followed by a Tukey-Kramer test as a post-hoc. For experiments where values didn’t follow the normal distribution, a non-parametric test was chosen: Kruskal-Wallis test followed by Dunn’s post hoc with Benjamini-Hochberg adjustment of p values. For each test the significance level α=0.05.

In plots, for intensity measurements, the individual values were displayed alongside means with ±95% confidence intervals. Line plots display only the mean values for each set of measurements and hence are accompanied with ribbon shading indicating standard deviation (s.d.). Information about the tests is given in figure legends.

## ACKNOWLEDGEMENTS

We thank the Bioimaging Center of the Faculty of Sciences at the University of Geneva, particularly Jerome Bosset for advice on imaging, and the Proteomic Facility in the Functional Genomics Center Zurich for proteomic data acquisition. We thank Michael Plank for outstanding advice on mass spectrometry and MS spectra interpretation, Dario Menendez and Kamila Delaney for some of the CRISPR injections, Alexandra Bondaz and Simona Abbatemarco for advice on spinning disc microscopy and data processing, Monica Gotta, Patrick Meraldi and Reto Gassmann for useful advice and Marina Berti for reagents preparation. We are grateful to past and present Steiner lab members for helpful discussions and to Monica Gotta, Patrick Meraldi and Isa Özdemir for comments on the manuscript. We are grateful to Gyorgyi Csankovszki for sharing the EKM36 strain and Reto Gassmann for sharing the GCP529 strain. Some strains were provided by the CGC, which is funded by the NIH Office of Research Infrastructure Programs (P40 OD010440). We thank the WormBase.

## AUTHOR CONTRIBUTION

JMW and FAS designed the study, JMW performed all genetic, staining, live imaging and genomic experiments, RFP carried out the IP-MS experiments, CGD performed protein purification and the *in vitro* kinase assays. JMW analysed the data. JMW and FAS wrote the manuscript with input from all authors. FAS obtained funding.

## DATA AVAILABILITY

All the data are contained within this manuscript and its supplements.

---

**Figure S1.**
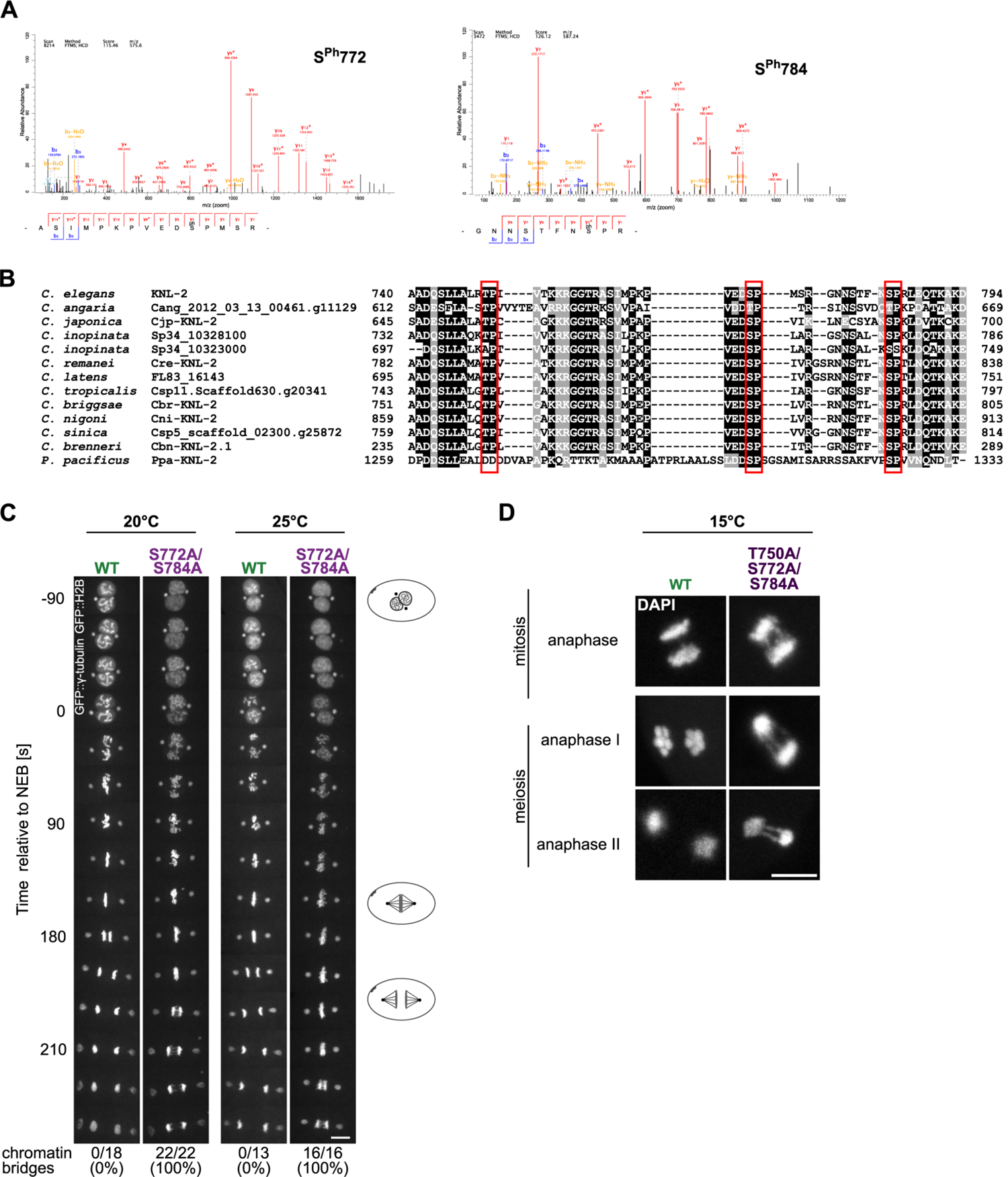
KNL-2 is phosphorylated on conserved residues T750, S772 and S784. (A) Exemplary spectra for peptides harbouring S772Ph or S784Ph modification identified by mass spectrometry. (B) Alignment of the C-terminal part of KNL-2 proteins from *Caenorhabditis* species, with KNL-2 from *Pristionchus pacificus* as an outgroup. The phosphosites identified in this study are marked with red boxes. (C) Kymographs illustrating the first embryonic division at 20°C and 25°C for wildtype (WT) and S772A/S784A embryos expressing GFP::H2B and GFP::γ-tubulin, with percentages of embryos exhibiting chromatin bridges during anaphase. NEB - nuclear envelope breakdown. Scale bar: 10 μm. (D) DAPI staining of fixed embryos from WT and T750A/S772A/S784A strains. Images show anaphases of mitotic and meiotic divisions at 15°C (permissive temperature for this strain). Scale bar: 5 μm.

**Figure S2.**
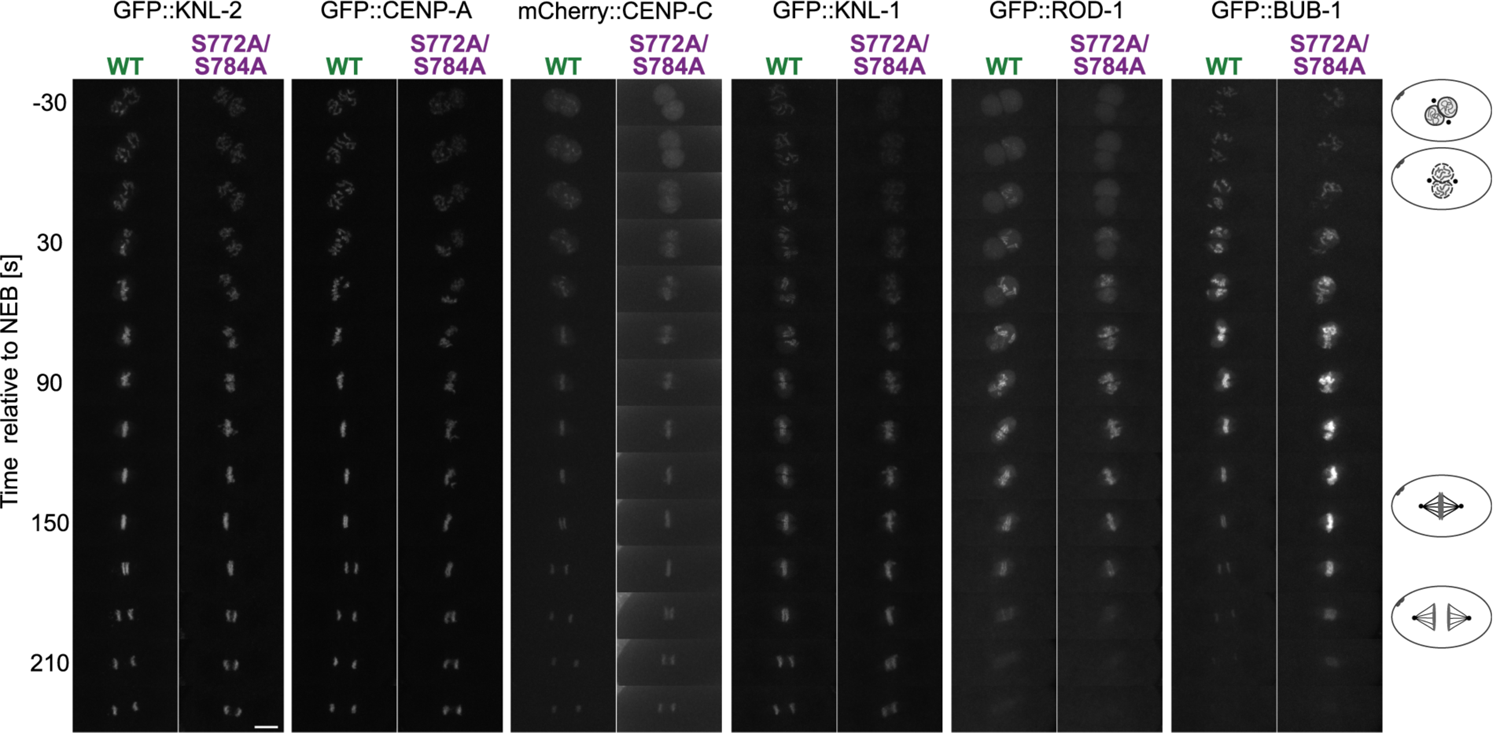
Localisation of centromeric and kinetochore proteins is not affected by S772A and S784A KNL-2 mutations. Kymographs showing the first embryonic division for wildtype (WT) and S772A/S784A embryos expressing the indicated GFP- or mCherry-tagged centromere and kinetochore proteins. NEB - nuclear envelope breakdown. Scale bar: 10 μm.

**Figure S3.**
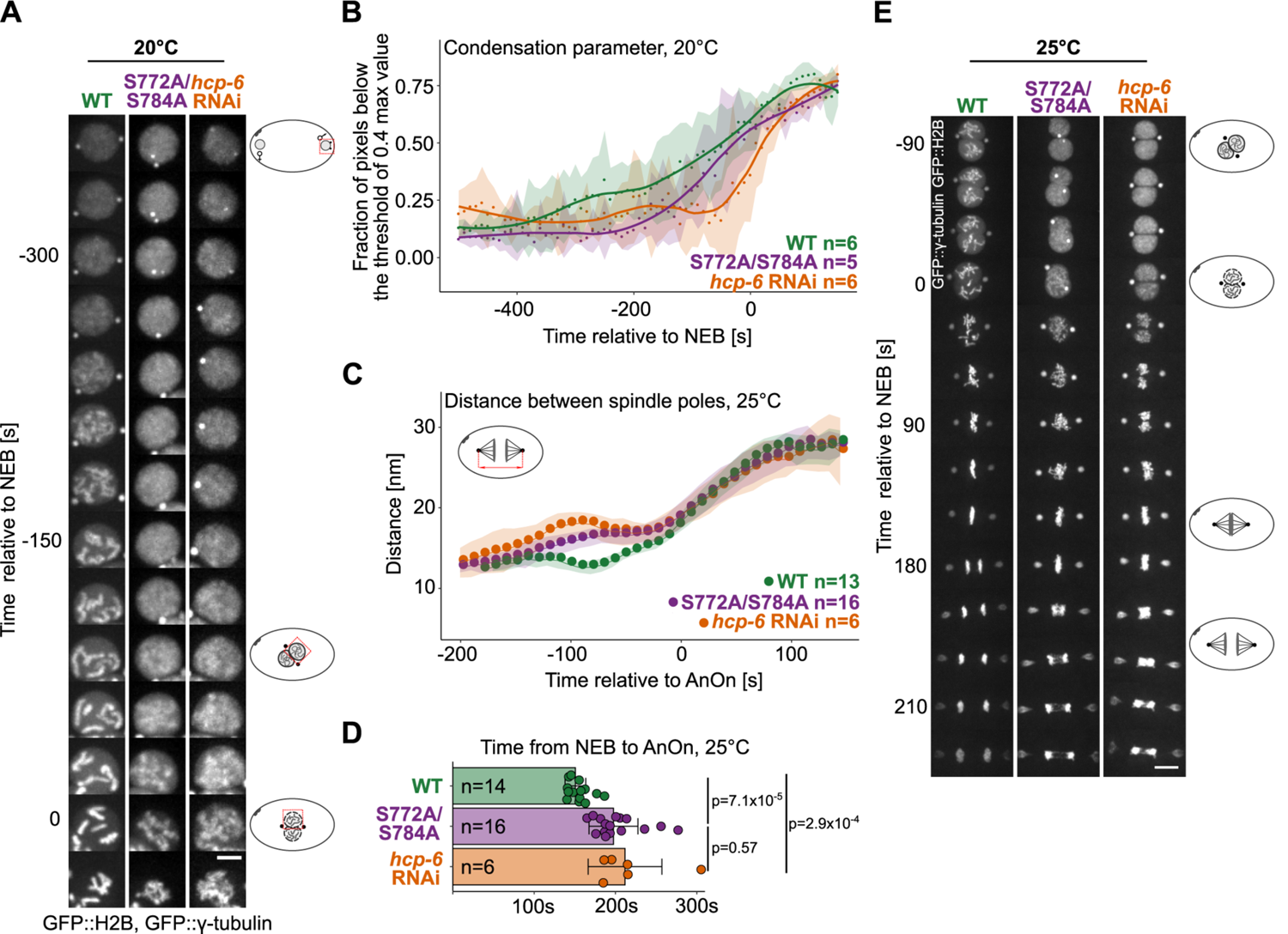
Condensation impairment and resulting phenotypes in the S772A/S784A strain. Wild type (WT), S772A/S784A and *hcp-6* RNAi strains expressing GFP::H2B and GFP::γ-tubulin were analysed. (A) Kymographs of male pronuclei, illustrating the progression of chromosome condensation over time at 20°C (permissive temperature). Scale bar: 5 μm. (B) Quantification of the condensation parameter at 20°C (fraction of pixels with values below the arbitrarily chosen threshold of 0.4 of the maximum pixel value) for the time series in (A). Dots show the mean value of the condensation parameter for each timepoint, shaded areas represent s.d. Line plots were fitted with the R loess function (span=0.4) for illustrating the trend. n corresponds to the number of embryos scored per condition. (C) Kymographs comparing the first embryonic division. NEB - nuclear envelope breakdown. Scale bar: 10 μm. (D) Graph illustrating the changes of the distance between spindle poles in time. Dots show the average distance for each timepoint, shaded areas represent s.d., n denotes the number of scored embryos. AnOn - anaphase onset. (E) Quantification of the time between NEB and AnOn. Dots correspond to individual embryos scored (the total number n is indicated), error bars depict s.d. Statistical significance was assessed with Kruskal-Wallis test followed by Dunn’s post hoc with Benjamini-Hochberg p-value adjustment.

**Figure S4.**
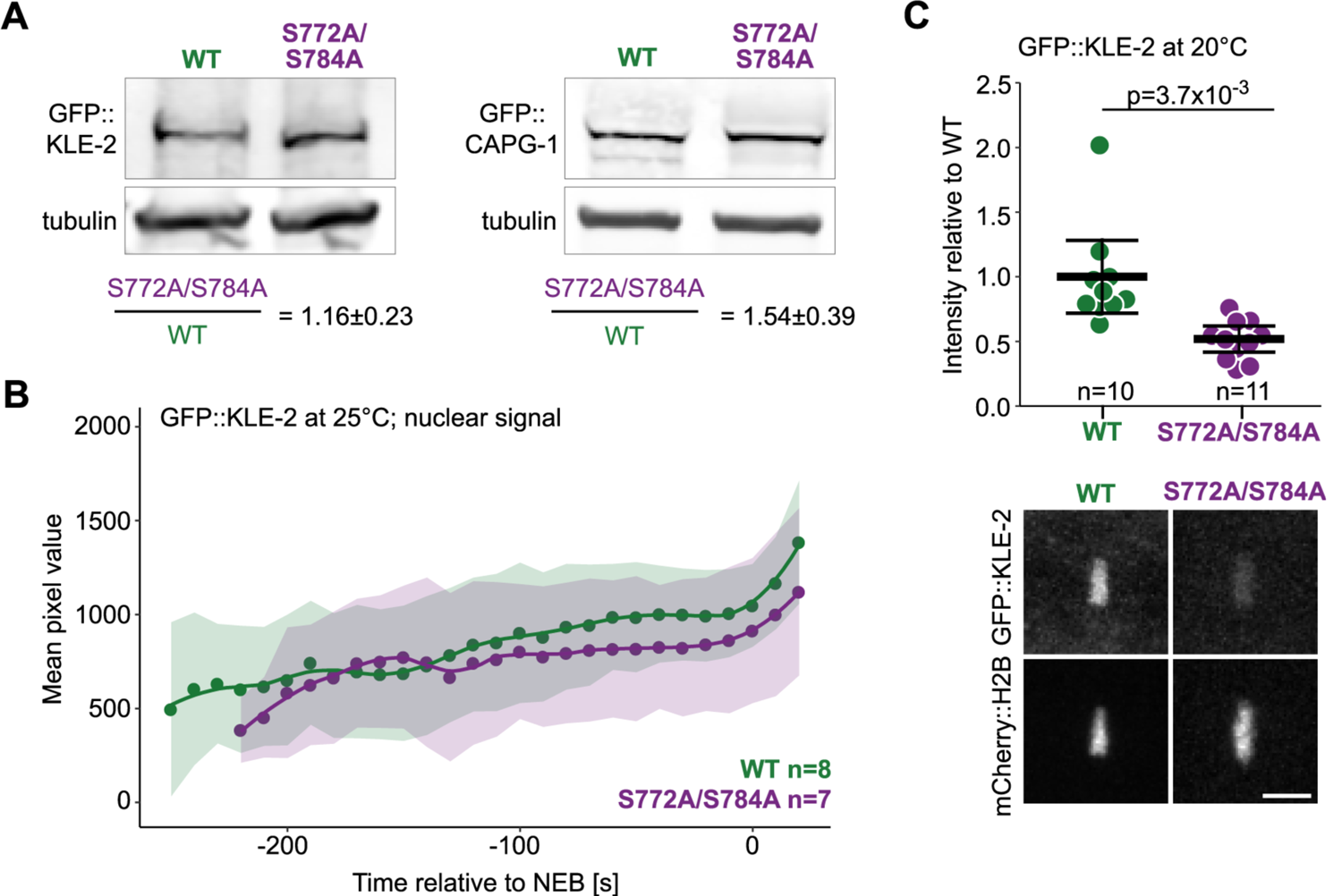
Condensin complexes in the S772A/S784A mutant. (A) Western blot of total embryonic lysates from wild type or S772A/S784A strains. An antibody against GFP was used for detecting GFP::KLE-2 or GFP::CAPG-1, tubulin was used as a loading control. GFP band intensities were normalised to tubulin band intensities, and condensin subunit abundance in the S772A/S784A strain was compared to wildtype (average and s.d. from three experiments). (B) Quantification of the nuclear GFP::KLE-2 abundance in one cell embryos of wildtype and S772A/S784A strains. The line plot represents average value for all measurements, shaded areas represent s.d., n corresponds to the number of scored embryos per condition. (C) Quantification of GFP::KLE-2 signal on first embryonic metaphase plates at 20°C (permissive temperature). Each data point represents one scored embryo. t-test was used for testing the statistical significance. Representative images are shown below the quantifications. Scale bar: 5 μm.

## SUPPLEMENTARY TABLES

**Table S1.** **Phosphopeptides identified in this study**

| Peptides | Modified residue no. | PSM | PEP | Localisation probability | Identified | Not phosphorylated peptide identified |
| --- | --- | --- | --- | --- | --- | --- |
| ASIMPKPVEDSP<br>MSR | 772 | 5 | 0.00016266<br>4 | 0.995874 | in 2<br>experiments | Yes |
| GNNSTFNSPR | 784 | 2 | 0.00024502<br>5 | 0.999924 | in 2<br>experiments | Yes |
| *TPIVTK* | *750* | *not identified in IP-MS experiments; T750 phosphorylated in in vitro assays* |  |  |  |  |
| Searches performed with MaxQuant (Version 1.6.0.16) |  |  |  |  |  |  |
| PSM - Peptide Spectrum Match; PEP - Posterior Error Probability; Localisation probability - denotes how likely a given residue is modified |  |  |  |  |  |  |

**Table S2.**
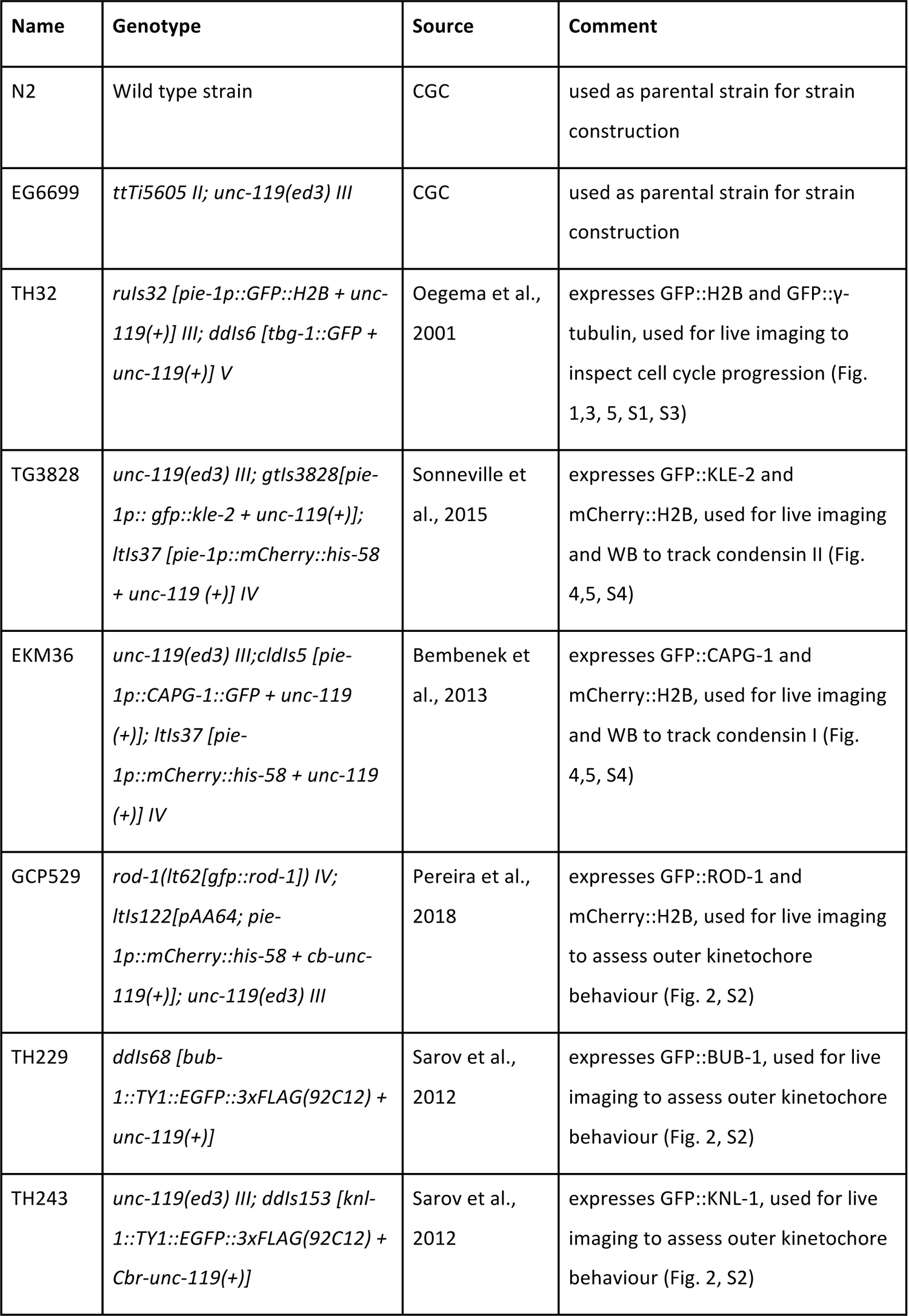
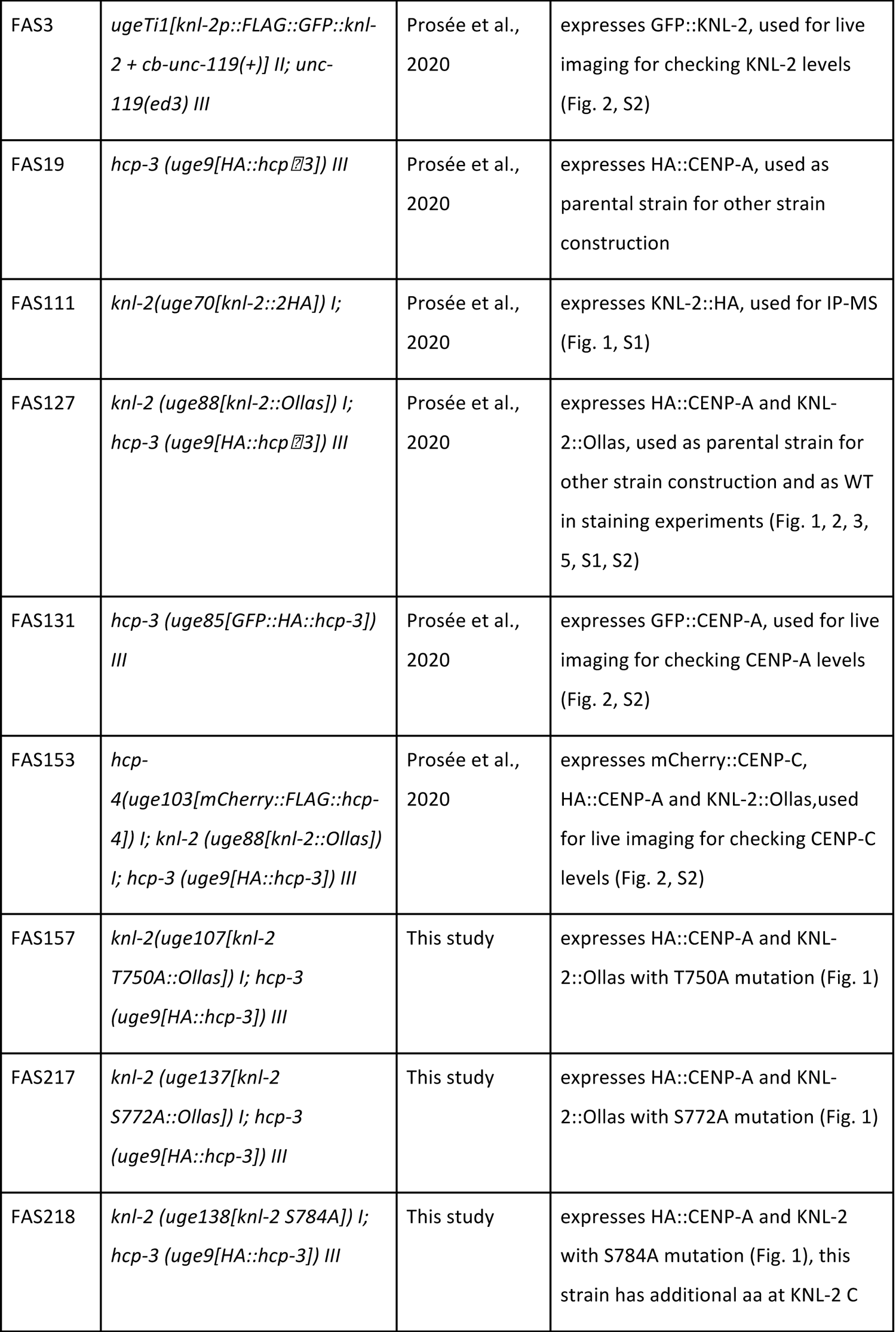
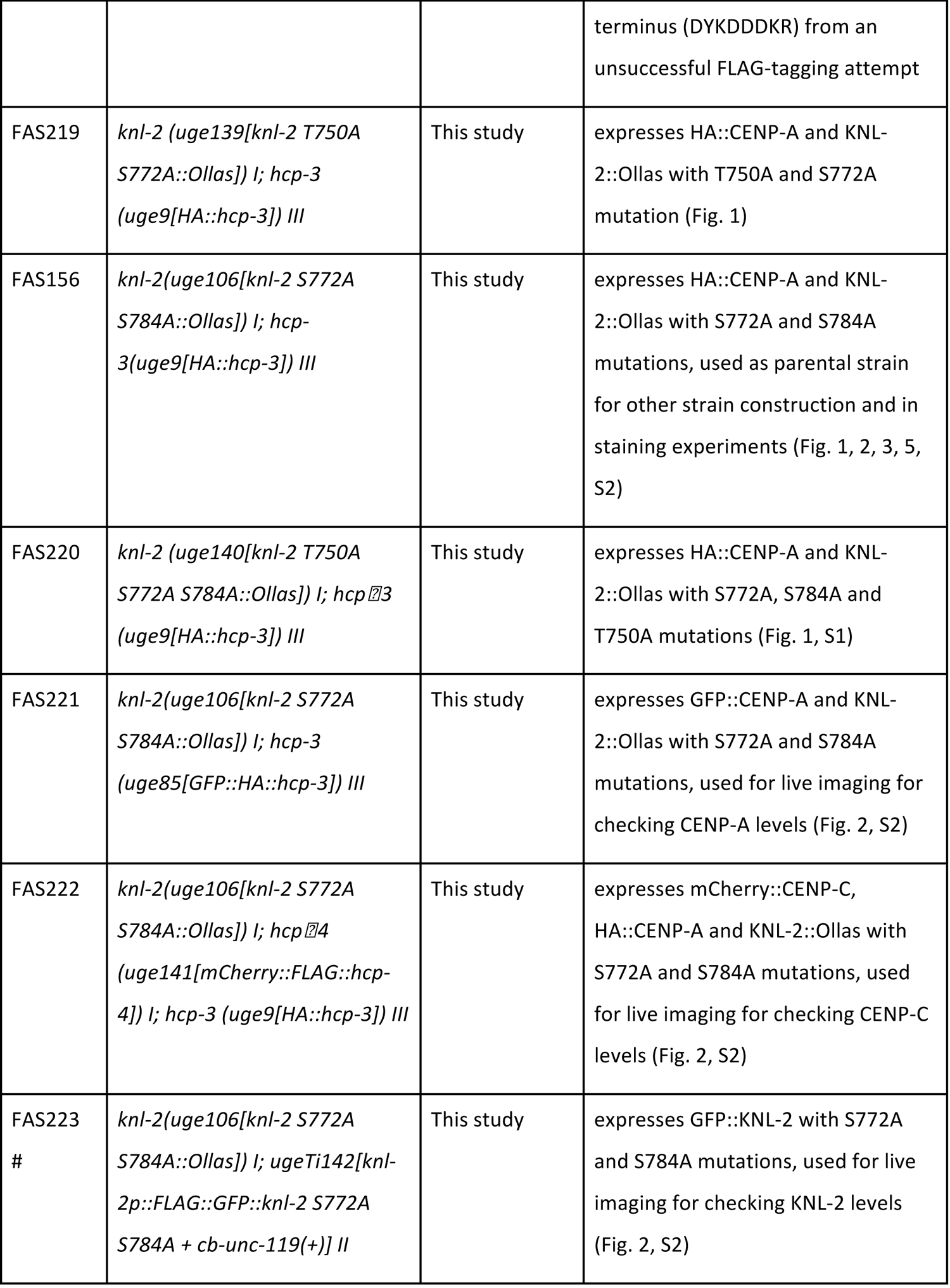
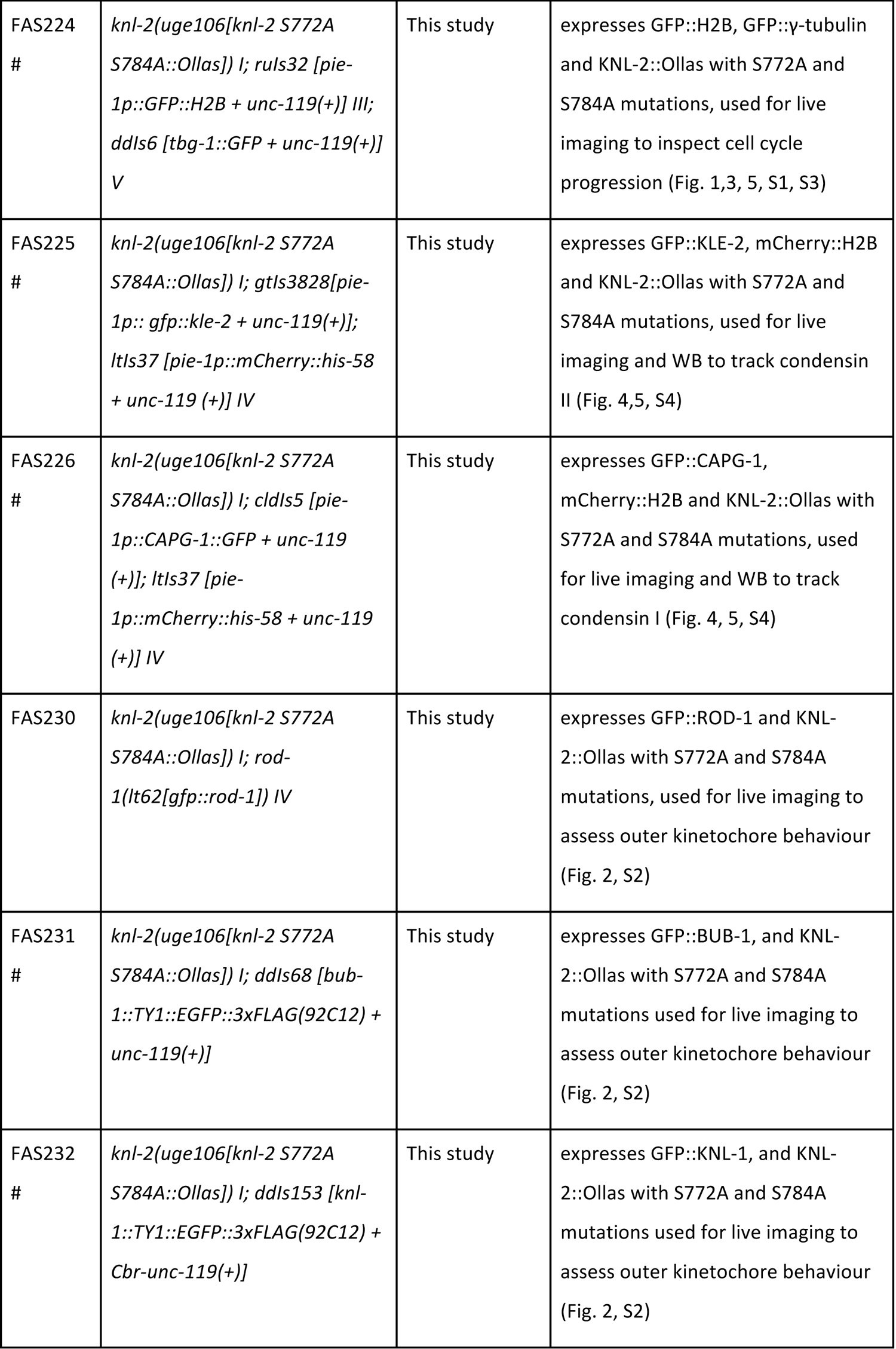
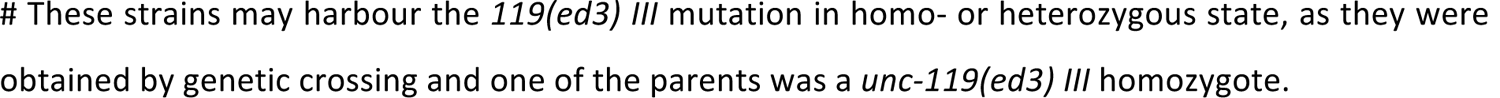
**List of strains used in this study**

**Table S3.**
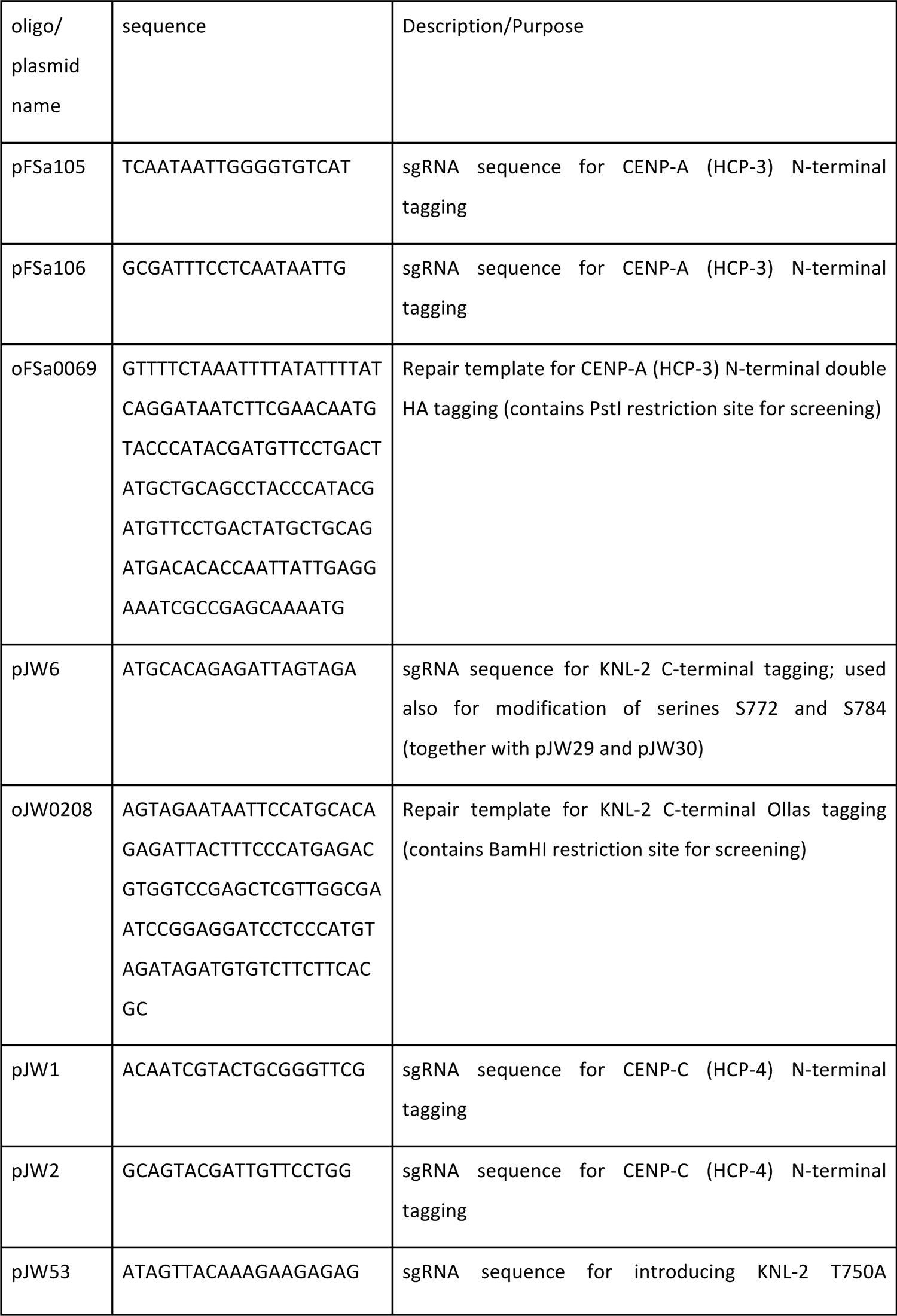
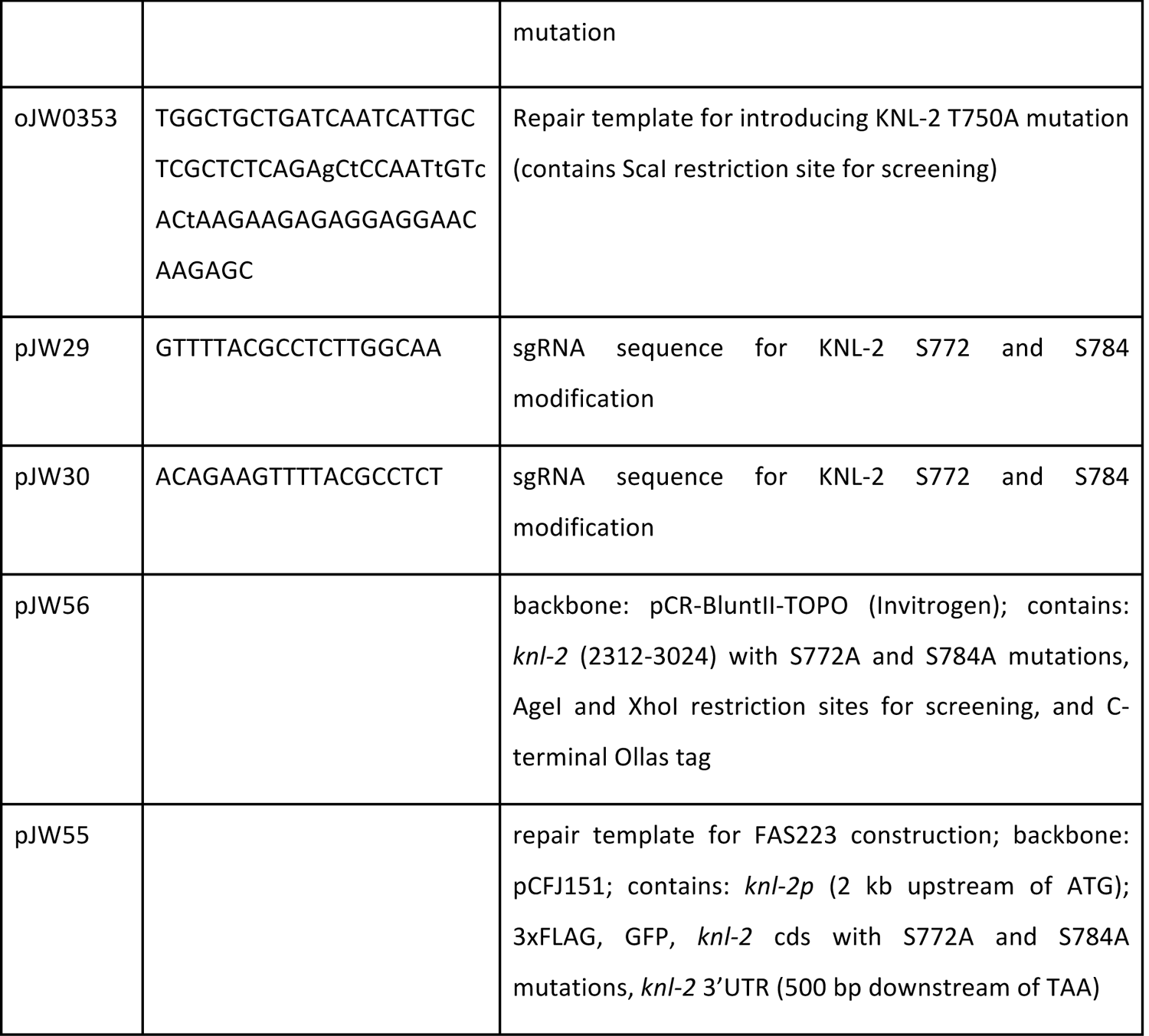
List of sgRNAs and repair templates used in this study

## MOVIES

Movie 1. First embryonic cell division in WT and the S772A/S784A strain expressing GFP::H2B and GFP::γ-tubulin at 20°C. Scale bar: 10 μm.

Movie 2. First embryonic cell division in WT and the S772A/S784A strain expressing GFP::H2B and GFP::γ-tubulin at 25°C. Scale bar: 10 μm.

Movie 3. First embryonic cell division in a strain expressing GFP::H2B and GFP::γ-tubulin at 25°C after partial *hcp-6* mRNA depletion. Scale bar: 10 μm.

Movie 4. First embryonic cell division in WT and the S772A/S784A strain expressing GFP::KLE-2 and mCherry::H2B at 25°C. Scale bar: 10 μm.

Movie 5. First embryonic cell division in WT and the S772A/S784A strain expressing GFP::CAPG-1 and mCherry::H2B at 25°C. Scale bar: 10 μm.

